# Spatial and Bulk Transcriptomic Profiling Defines the Molecular Evolution of Cutaneous Squamous Cell Carcinoma and Reveals Stage-Specific Biomarkers of Clinical Relevance

**DOI:** 10.64898/2026.04.30.721943

**Authors:** Fatima Naji, Sergio Oterino-Sogo, Fanny Beltzung, Daniel García-Ruano, Walid Mahfouf, Jean-Philippe Guegan, Mylene Bohec, Alexis Groppi, Marie Beylot-Barry, Léa Dousset, Macha Nikolski, Hamid-Reza Rezvani

## Abstract

Cutaneous squamous cell carcinoma (cSCC) is a common skin cancer associated with substantial morbidity and mortality in advanced stages. Despite its well-described stepwise progression from actinic keratosis to invasive disease, robust molecular markers for stage discrimination and clinical decision-making remain limited. We sought to define the transcriptional continuum underlying cSCC progression, identify stage-associated biomarkers, and assess the broader relevance of these programs across human malignancies.

Bulk RNA sequencing (HTG EdgeSeq) and spatial transcriptomics (GeoMx) were performed on biopsies from eight patients, each presenting multiple disease stages (healthy skin, premalignant lesion, tumor core, and invasive front) within the same lesion field, enabling within-patient analysis of progression.

Spatial transcriptomic analyses identified more than 2,000 differentially expressed genes whose expression varied across disease stages. These genes were organized into 18 coordinated expression programs reflecting progressive biological rewiring during tumor evolution. Proliferation, extracellular matrix remodeling, inflammation, and stress-response pathways were progressively upregulated, whereas epithelial differentiation and metabolic processes, including lipid and amino acid metabolism, were downregulated. Macrophages exhibited distinct metabolic reprogramming, with increased purine metabolism, glycolysis, and pyruvate metabolism across progression.

To evaluate the broader clinical relevance of these progression-associated programs, we developed a reproducible Snakemake pipeline to systematically screen 32 solid and hematologic malignancies from The Cancer Genome Atlas (TCGA). A combined cSCC-progression signature was significantly associated with poor overall survival (*P* < 0.05) in 10 additional cancer types. Finally, we identified 12 stage-informative biomarkers, whose spatially restricted expression patterns were validated using Visium HD.

This study provides a spatially resolved and stage-aware transcriptomic map of cSCC progression, identifies coordinated gene programs underlying disease evolution, and defines progression-associated signatures with prognostic relevance across multiple cancers, highlighting their potential translational value.

## Introduction

Cutaneous squamous cell carcinoma (cSCC) is the second most common keratinocyte carcinoma after basal cell carcinoma (BCC) ^1,2^. Compared with BCC, cSCC exhibits greater clinical aggressiveness, with a measurable risk of metastasis ^3^, disease-specific mortality^4^, and substantial morbidity ^5^. It arises from the malignant transformation of epidermal keratinocytes ^6^, primarily driven by cumulative ultraviolet (UV) exposure ^7^ and its incidence continues to increase worldwide due to aging populations and chronic UV exposure ^1,2^.

cSCC evolves along a multistep continuum, progressing from precancerous actinic keratosis (AK) through *in situ* cSCC and ultimately to invasive disease ^8^. Although diagnosis and management rely on clinical and histopathological criteria, current grading and risk-stratification systems ^9–12^, remain insufficient to accurately predict progression across this continuum ^1314^. This underscores two unmet needs: robust biomarkers to molecularly stage cSCC and refine risk assessment, and a deeper understanding of the transcriptional programs governing stage transitions, including tumor-intrinsic and microenvironmental contributions ^15^.

Transcriptomic technologies have expanded insight into cSCC biology ^16,17^. Bulk RNA sequencing studies enabled large-cohort analyses of progression-associated changes ^17^, while single-cell RNA sequencing revealed cell type-specific alterations but lacked spatial context and was often limited by sample sizes ^16^. Spatial transcriptomics ^18^ now permits *in situ* gene expression profiling within preserved tissue architecture, uncovering microenvironmental differences between invasive and premalignant lesions ^19,20^ and characterizing tumor–stroma interfaces ^19^. However, most studies have examined isolated disease stages or inferred progression across different patients, leaving the full primary cSCC progression continuum incompletely defined ^19,20^.

Here, we address this gap by analyzing patient samples that encompass multiple pathological stages within the same field of cancerization, enabling reconstruction of coherent progression-associated gene programs across the cSCC continuum. We combined complementary bulk and spatial transcriptomic approaches and then assess the broader clinical relevance of identified cSCC progression-associated gene signatures across multiple cancer types. Key findings were further validated using Visium HD spatial transcriptomics in additional continuum cases, providing cross-platform confirmation within the same lesion field.

## Materials and Methods

### Cohort of Patients

The MITOSKIN project is a prospective translational study conducted in the Department of Dermatology, Bordeaux University Hospital. Patient presenting with lesions clinically suspected to represent different stages of cSCC were enrolled after written informed consent. For each patient, one biopsy was formalin-fixed for histopathologic confirmation and one was snap-frozen for molecular analyses. An additional snap-frozen biopsy of adjacent healthy skin (≥5 cm from the tumor margin) was collected for comparison.

For this study, eight patients exhibiting multiple lesion stages within the same field of cancerization (healthy skin, AK/ cSCC *in situ*, invasive cSCC, and invasive front) were selected to enable within-patient reconstruction of disease progression. Clinical and pathological characteristics are provided in **Supplementary Table S1**.

All procedures were conducted in accordance with national regulations governing biomedical research involving human participants. Written informed consent was obtained from all participants. Ethical approval was granted by the French National Commission for Data Protection and Liberties (approval ID-RCB 2018-A03096; January 4, 2019) and the study was registered at ClinicalTrials.gov (NCT04389112).

Additional materials and methods are provided in additional supplementary material file (**Appendix S1**).

## Results

### Bulk transcriptomic profiling by HTG EdgeSeq defines global stage-associated gene expression trends in cSCC

To minimize inter-individual variability and reconstruct coherent stage-associated transcriptional programs, we analyzed a cohort of eight patients presenting multiple stages of the cSCC continuum within the same field of cancerization, including adjacent healthy skin, actinic keratosis/cSCC in situ, invasive tumor core, and invasive front (**Figure 1A**). Such specimens are relatively rare in clinical practice and enable a within-patient design that more accurately captures progression-associated molecular changes. Because AK and cSCC *in situ* could not always be reliably distinguished histologically when displaying overlapping features, these lesions were analyzed as a single category to reduce potential misclassification. Leveraging this unique cohort, we combined bulk RNA sequencing (HTG EdgeSeq; HTG-Seq) with spatial transcriptomics (GeoMx Digital Spatial Profiler; GeoMx) to define transcriptional programs associated with cSCC progression (**Figure 1B; Supplementary Figures S1A-B**).

**Figure 1.**
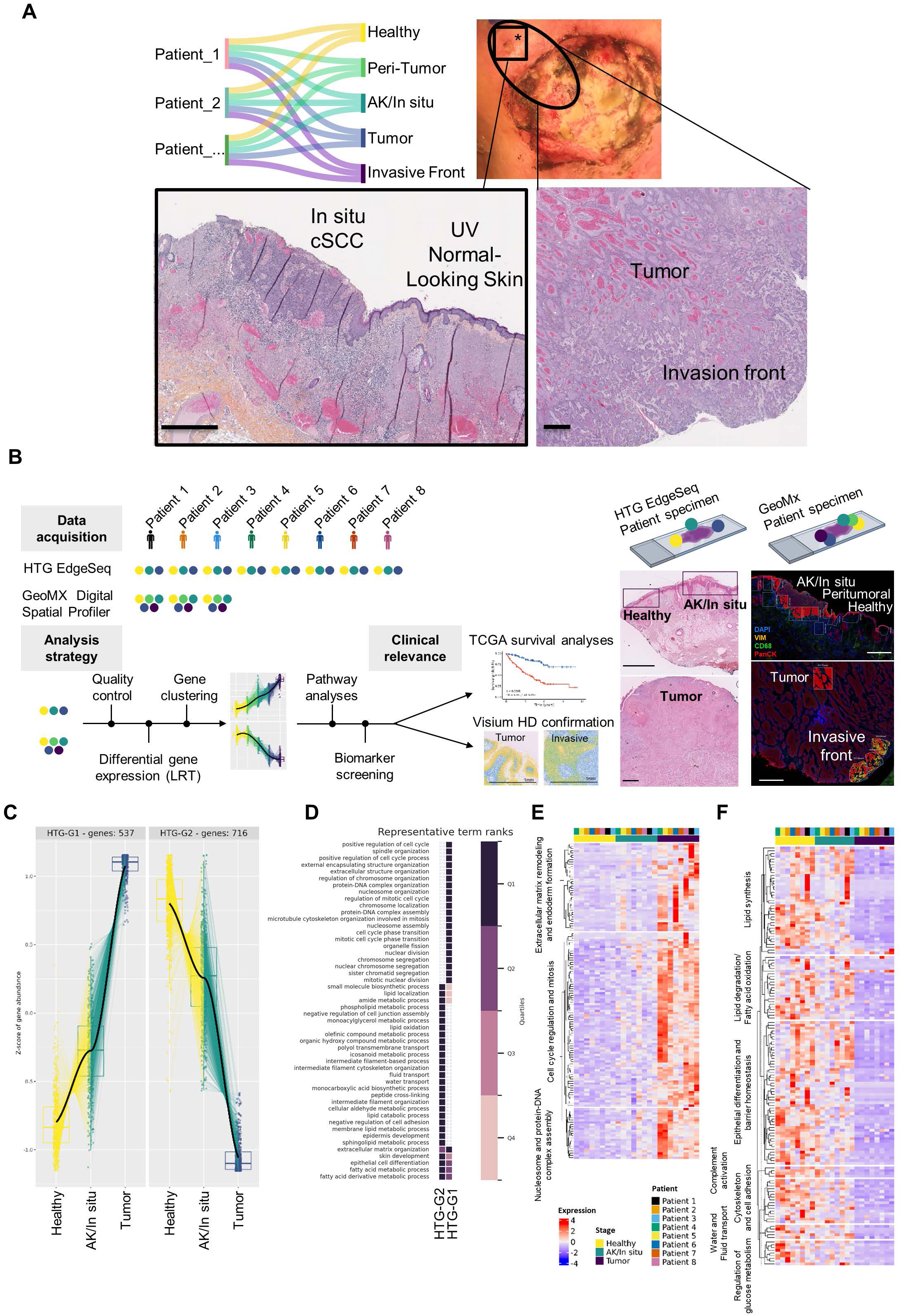
Bulk transcriptomic profiling by HTG EdgeSeq identifies two stage-associated gene expression trends during cSCC progression. **A.** Patients presenting multiple stages within the same field of cancerization, including healthy skin, actinic keratosis/cSCC in situ, invasive tumor core, and invasive front were included in this study (right panel). A representative case is shown, in which both *in situ* cSCC and invasive cSCC are present within a single lesion field. Hematoxylin and eosin (H&E) staining illustrates the spatial identification of distinct stages within one biopsy, including adjacent healthy skin, peritumoral region, cSCC *in situ*, invasive tumor core, and invasive front. **B.** Schematic overview of the study design and analyses. Two transcriptomic technologies were used: HTG EdgeSeq (HTG-Seq) and NanoString GeoMx Digital Spatial Profiler (DSP). Representative H&E-stained images illustrate identification of multiple disease stages within the same biopsy from individual patients. For GeoMx analyses, tissue sections were stained with a CD68/PanCK/Vimentin/Sytox multiplex panel to visualize tissue architecture and guide selection of regions of interest (ROIs). CD68⁺ corresponded to macrophages, PanCK⁺ to epithelial cells, and PanCK⁺/Vimentin⁺ regions to the invasive tumor front. Scale bar, 1 mm **C.** Likelihood ratio test (LRT) applied to HTG-Seq data identified two major groups (HTG-G1, HTG-G2) of differentially expressed genes (DEGs) exhibiting opposite, stage-dependent expression trajectories (progressively upregulated versus progressively downregulated) across cSCC progression. **D.** Gene Ontology Biological Process (GO: BP) pathway enrichment analysis of DEGs in HTG-G1 and HTG-G2. Following ORSUM summarization, top representative GO:BP terms are shown, with quartile ranksindicated. **E-F.** Heatmaps displaying DEGs and associated biological pathways that are progressively upregulated in HTG-G1 (**E**) or progressively downregulated in HTG-G2 (**F**) across healthy skin, AK, and tumor stages. The lists of genes found to be differentially expressed and the gene lists used for the heatmaps can be found in https://services.cbib.u-bordeaux.fr/cSCC_gene_tables/.

HTG-Seq profiling was performed on healthy skin, AK/*in situ*, and invasive tumor regions from all eight patients (**Figure 1B**). Progression-associated genes were identified using a likelihood ratio test (LRT) with disease stage as the model factor, enabling simultaneous comparison across all stages while reducing false positives inherent to multiple pairwise contrasts. Differentially expressed genes (adjusted *P* < 0.01) were clustered, revealing two principal gene expression trajectories: 537 genes progressively upregulated (HTG-G1) and 716 genes progressively downregulated (HTG-G2) across cSCC progression (**Figure 1C**). To functionally interpret these expression patterns, pathway enrichment analysis was performed using g:Profiler against Gene Ontology Biological Process (GO:BP) terms and related gene sets. Redundant terms were summarized using ORSUM, and representative enriched processes are shown in **Figure 1D**. Progression-upregulated genes (HTG-G1) were significantly enriched for cell cycle regulation, mitosis, nucleosome assembly, and extracellular matrix (ECM) remodeling (**Figure 1E**). In contrast, progression-downregulated (HTG-G2) were associated with epithelial differentiation, lipid and glucose metabolism, cytoskeletal organization, and aquaporin-mediated transport (**Figure 1F**).

Together, these results define broad coordinated transcriptional programs that dynamically evolve along the cSCC progression continuum.

### Spatial transcriptomics identifies 18 gene expression patterns associated with cSCC progression and specific stages

To refine bulk-derived signatures, we performed spatial transcriptomic profiling using the GeoMx, enabling high-resolution characterization of stage-specific regions across the cSCC continuum. Three patients exhibiting a continuous histopathological spectrum (healthy skin, AK/cSCC *in situ*, invasive tumor core, and invasive front) were included (**Figure 1B; Supplementary Figures S2-S3**). Guided by multiplex immunofluorescence–based tissue segmentation, 43 areas of illumination (AOIs) were profiled across defined epithelial PanCK^+^) and macrophage-enriched (CD68^+^) compartments spanning disease stages (**Supplementary Figure S2-S3**; **Supplementary Table S2**).

Applying LRT within PanCK^+^ compartments, we identified 18 distinct gene groups (Ker-G1 to Ker-G18) displaying progression-associated or stage-dependent expression patterns across the continuum of cSCC (**Figure 2A**). These spatially resolved expression trends provided a structured framework for interrogating the biological pathways underpinning cSCC evolution. Based on expression dynamics and biological relevance, gene groups with robust and consistent patterns were selected and consolidated into three principal classes: (i) progressively upregulated groups (groups Ker-G1 and -G3), (ii) progressively downregulated groups (groups Ker-G2 and -G4), and (iii) metabolically associated groups (groups Ker-G2, -G4, -G6, and -G10). Representative GO:BP terms for these selected groups are summarized in **Figure 2A-B**.

**Figure 2.**
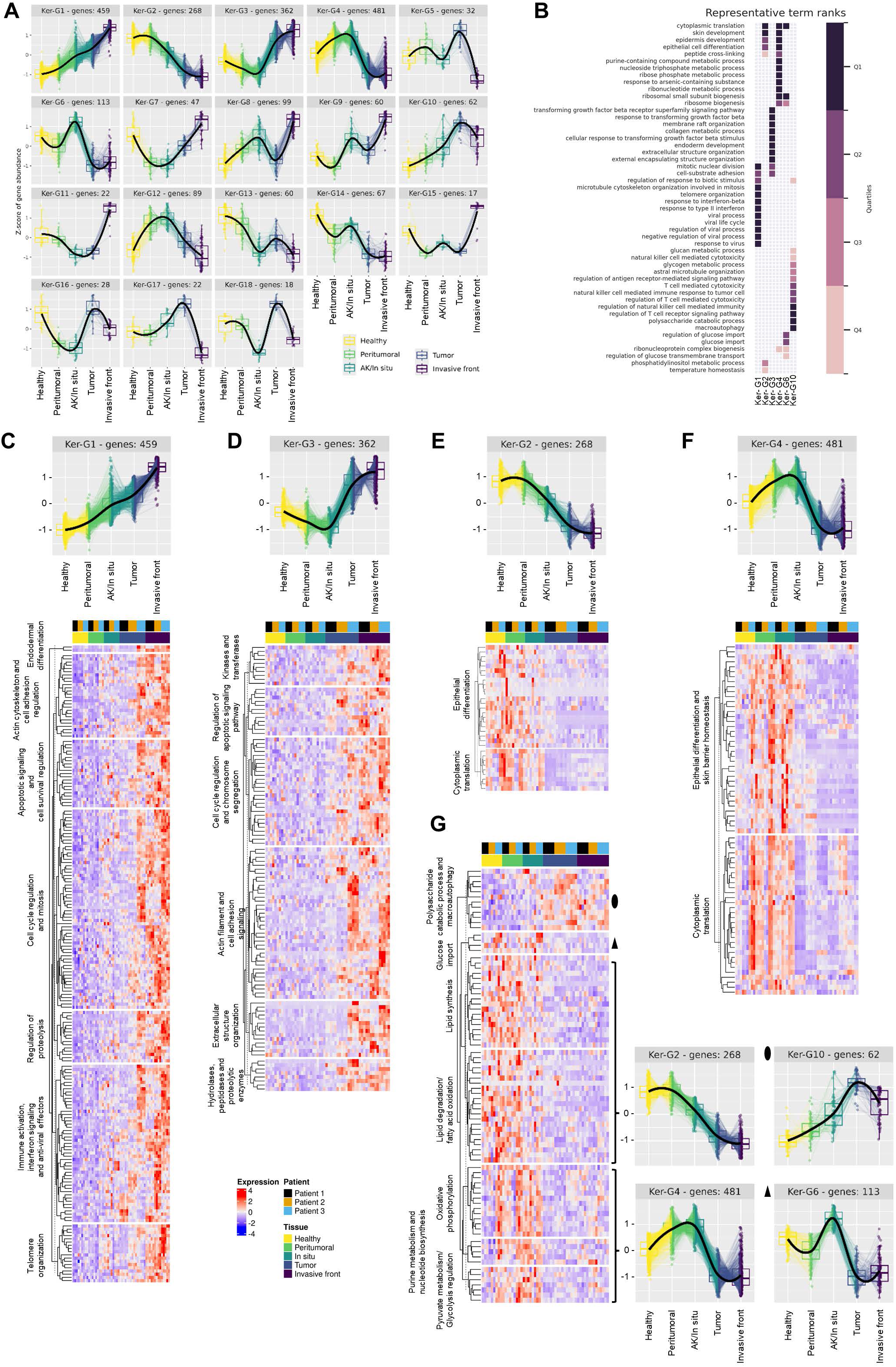
Spatial transcriptomic analysis of PanCK⁺ regions reveals 18 distinct cSCC progression and stage-associated transcriptional programs. **A.** LRT applied to PanCK⁺ regions in the GeoMx dataset identified 18 gene expression trajectories representing distinct stage-dependent trends of DEGs across cSCC progression. **B.** GO:BP pathway analysis of DEGs from selected trajectories, including upregulated (gene groups Ker-G1 and -G3), downregulated (gene groups 2 and 4), and metabolically reprogrammed trends (gene groups Ker-G2, Ker-G4, Ker-G6, and Ker-G10). Following ORSUM summarization, top representative GO:BP terms are shown, with rank quartiles indicated. **C-D.** Heatmaps of DEGs from upregulated gene groups Ker-G1 (n = 459 genes) and Ker-G3 (n = 362 genes), showing progressive increases in expression across healthy skin, peritumoral regions, AK/*in situ*, tumor, and invasive front. Enriched pathways include immune and interferon signaling, cell cycle regulation, cytoskeletal organization, cell adhesion, and extracellular matrix organization. **E-F.** Heatmaps of DEGs from downregulated gene groups Ker-G2 (n = 268 genes) and Ker-G4 (n = 481 genes), showing progressive decreases in expression across cSCC stages. Enriched pathways include cytoplasmic translation, epithelial differentiation and skin development. **G.** Heatmap of gene groups reflecting metabolic reprogramming during cSCC progression (gene groups Ker-G2, -G4, -G6, and -G10). These trajectories demonstrate complex, stage-dependent alterations in metabolic pathways, including downregulation of lipid metabolism, dynamic regulation of glycolysis and oxidative phosphorylation, activation of autophagy-related processes (glycophagy and mitophagy), and modulation of glucose import, consistent with shifts between anabolic and catabolic metabolic states. The order of the genes in the heatmaps reflects that of **Supplementary Materials**.

Among the upregulated gene groups, Ker-G1 (459 genes) demonstrated sustained upregulation from healthy skin through invasive carcinoma and was enriched for immune and interferon signaling, cell cycle progression and mitosis, cytoskeleton organization and cell adhesion, apoptosis regulation, and proteolysis (**Figure 2C**), alongside markers of altered epithelial differentiation (e.g., Laminin subunit alpha-3 and beta-3 (*LAMB3*, *LAMA3*)). Ker-G3 (362 genes) exhibited modest early reduction followed by progressive upregulation in later disease stages. While this group shared several functional themes with Ker-G1 (such as enhanced cytoskeletal dynamics, cell adhesion, and regulation of cell cycle and apoptotic processes), it was distinguished by strong enrichment in ECM organization and remodeling pathways, including collagen metabolism and mesenchymal transition–related processes (**Figure 2D**). Together, these data suggest that Ker-G1 reflects early tumor-intrinsic immune activation, whereas Ker-G3 captures later ECM remodeling and invasive programs, providing temporal resolution of cSCC progression. In contrast, downregulated gene groups reflected progressive loss of epithelial identity and biosynthetic capacity. Ker-G2 (268 genes) showed continuous decline across stages and was enriched for processes related to cytoplasmic translation machinery, including ribosomal function and protein synthesis, as well as epithelial differentiation, skin development, and lipid metabolism pathways (**Figure 2E**). Ker-G4 (481 genes) displayed transient early upregulation followed by sustained downregulation in later advanced stages. In addition to overlapping differentiation and translational signatures with Ker-G2, this group uniquely encompassed oxidative phosphorylation, purine metabolism, nucleotide biosynthesis, and regulation of glycolysis (**Figure 2F**). Collectively, these groups indicate progressive attenuation of epithelial differentiation accompanied by a broad reorganization of metabolic and biosynthetic processes during cSCC evolution.

Focusing on metabolic pathways, four gene groups (Ker-G2, Ker-G4, Ker-G6, and Ker-G10) were particularly associated with metabolic reprogramming during cSCC progression (**Figure 2G**). Lipid metabolism pathways were consistently downregulated, whereas glycolysis and oxidative phosphorylation displayed increased at early stages (healthy skin to AK/*in situ*) and declined in advanced disease. Ker-G10 was enriched for autophagy-related processes, including mitophagy, glycophagy, and polysaccharide catabolism, consistent with adaptive responses to metabolic stress. Meanwhile, insulin signaling and glucose import pathways represented in Ker-G6 showed early upregulation followed by reduction. Together, these findings suggest that early cSCC retains greater metabolic flexibility, whereas advanced stages display transcriptional signatures consistent with stress adaptation and energy conservation.

### Spatial transcriptomics reveal opposing metabolic reprogramming in macrophages compared to tumor cells during cSCC progression

To examine immune microenvironment dynamics, we profiled CD68⁺ areas, corresponding to macrophage (Mac) compartments. Likelihood ratio testing identified two principal trajectories: progressive upregulation (Mac-G1) and downregulation (Mac-G2) across cSCC stages (**Figure 3A**), consistent with stage-dependent immune remodeling.

**Figure 3.**
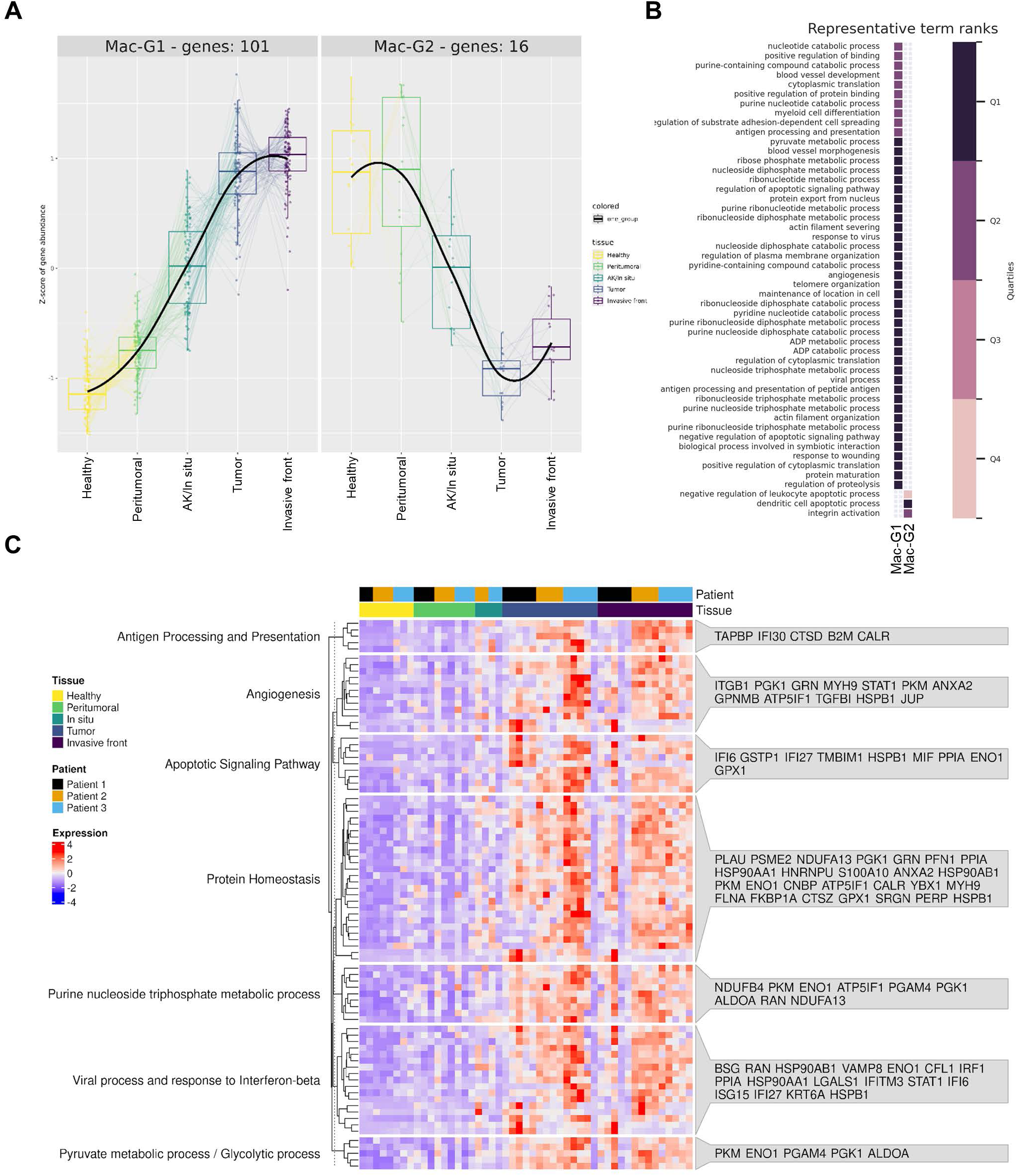
Spatial transcriptomic analysis of CD68^+^ regions reveals progressive immune activation and metabolic remodeling. **A.** LRT applied to macrophages-enriched (CD68+) regions in the GeoMx dataset. Two opposite, stage-dependent expression trends (progressively upregulated and progressively downregulated) were identified across cSCC progression. **B.** GO: BP pathway analysis of DEGs in the upregulated and downregulated trajectories. Following ORSUM summarization, top representative GO:BP terms are shown, with rank quartiles indicated. **C.** Heatmap shows expression patterns of DEGs of group Mac-G1 (n = 101 genes) from CD68+ macrophage-enriched regions across cSCC progression. Pathway analysis identified five main categories: interferon signaling and viral response, purine and glycolytic metabolism, protein homeostasis, apoptotic signaling regulation, and angiogenesis. Interestingly, metabolic pathways such as glycolysis, downregulated in tumor cells, were upregulated in macrophages, underscoring divergent tumor–immune metabolic adaptations. The order of the genes in the heatmaps reflects that of **Supplementary Materials**.

Pathway enrichment analysis and ORSUM summarization achieved significant results predominantly within Mac-G1 (101 genes), which showed sustained upregulation from early to advanced lesions (**Figure 3B**). Enriched processes included interferon signaling, antiviral responses, and antigen presentation pathways (**Figure 3C**), reflecting macrophage activation within the tumor microenvironment. Notably, macrophages demonstrated upregulation of purine metabolism, glycolysis, and pyruvate metabolism, patterns contrasting sharply with the progressive downregulation of these same pathways observed in PanCK⁺ tumor cells. These data indicate divergent metabolic adaptation, with tumor cells adopting a restrained phenotype while macrophages increase metabolic activity during progression. Additional enriched processes included protein homeostasis and remodeling, regulation of apoptotic signaling, and angiogenesis, suggesting coordinated metabolic and functional adaptation of macrophages during tumor progression.

Comparison with bulk HTG-Seq signatures highlights the complementary value of spatial profiling. Although both approaches identified shared progression-associated programs (including increased cell cycle and EMC remodeling pathways and decreased lipid metabolism and epithelial differentiation), immune and interferon signaling programs detected in spatially defined tumor regions were largely absent in bulk data, underscoring how whole-tissue analyses can obscure cell-type–specific transcriptional programs.

### A spatially identified cSCC-progression signature correlates with poor disease overall survival in multiple different human cancers

To assess the generalizability of the molecular programs associated with cSCC progression, we analyzed RNA-seq data from 32 cancer types available through The Cancer Genome Atlas (TCGA). To ensure reproducibility and facilitate the reusability of this analysis framework, we developed a Snakemake pipeline that automates pan-cancer screening of gene- and gene set–based survival associations **(Figure 4A**, see methods). We next applied this pipeline to the GeoMx-derived gene sets upregulated (see group Ker-G1; **Figure 2C**) and downregulated (see group Ker-G2; **Figure 2E**) during cSCC progression. These gene sets were integrated into a unified cSCC-progression signature, which was used to stratify cancer patients. The 25^th^ percentile of cases with the highest and lowest enrichment for each cohort were screened for overall survival. This signature significantly separated patient survival outcomes in 12 of the 32 cancer cohorts (log-rank *P* < 0.05; **Figure 4B**). In 10 of these cohorts, high signature scores were associated with poor overall survival, indicating that cSCC-linked transcriptional programs broadly reflect aggressive tumor behavior across diverse malignancies. Notably, ovarian cancer and skin cutaneous melanoma represented exceptions, in which high signature scores correlated with improved survival (**Figure 4C**), suggesting context-dependent biological effects.

**Figure 4.**
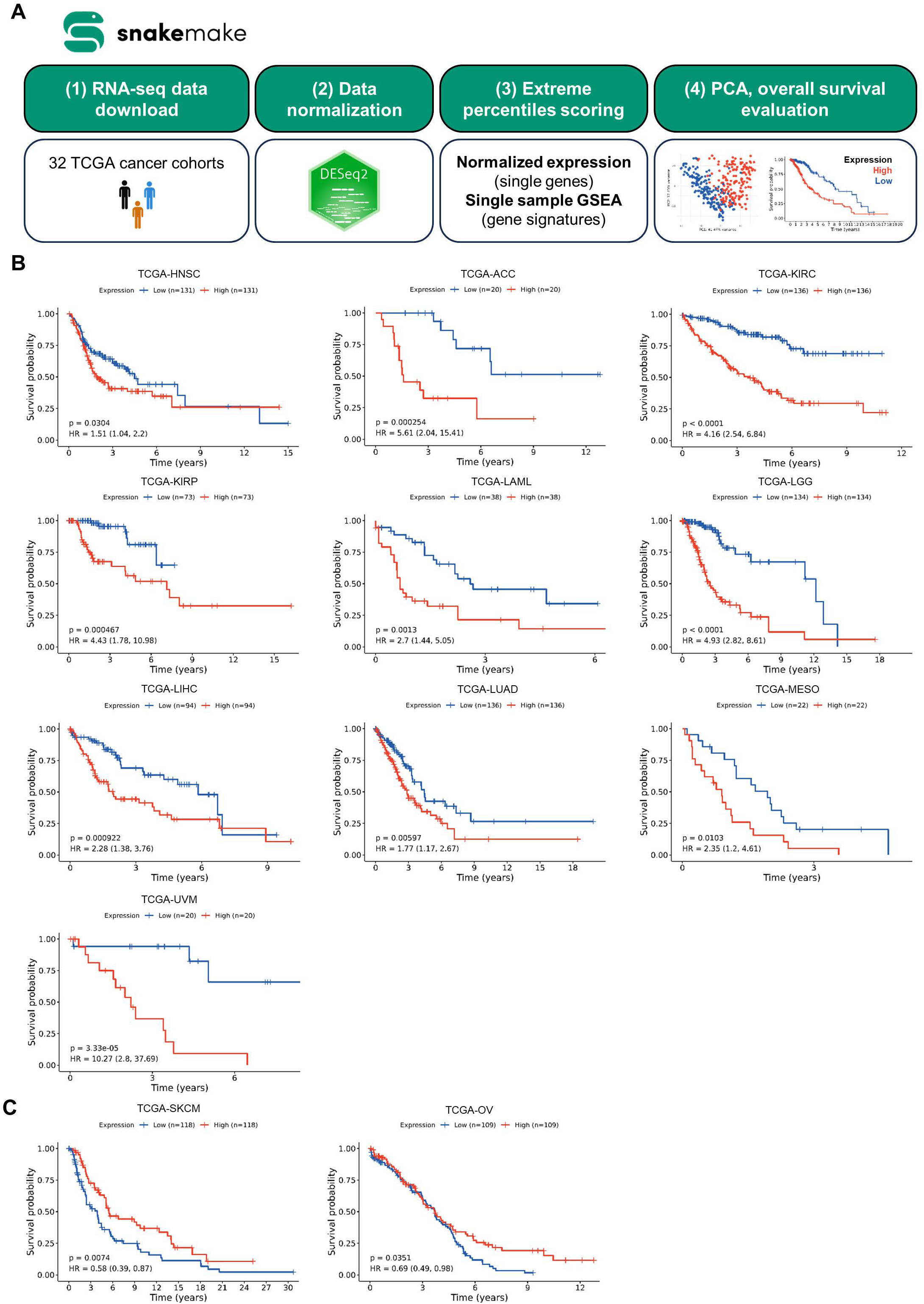
A cSCC-progression molecular signature correlates with poor overall survival across multiple TCGA cancer cohorts. **A.** Schematic of the Snakemake pipeline used to screen The Cancer Genome Atlas cohorts. GeoMx-derived gene sets identified as upregulated (Ker-G1) and downregulated (Ker-G2) during cSCC progression were used to compute single-sample gene set enrichment analysis (ssGSEA) scores for each cohort. The pipeline integrates upregulated and downregulated gene sets by calculating a combined progression score (Ker-G1 score minus Ker-G2 score), enabling robust application of spatially derived differential expression signatures to external datasets. Patients were stratified into high- and low-score groups using median (25th percentile) cutoffs for each cancer type. **B.** Pan-cancer interrogation of TCGA cohorts using the combined cSCC-progression signature. Median-based stratification identified significantly worse overall survival in 10 of 32 cancer cohorts (log-rank *P* < 0.05), indicating that cSCC progression–associated transcriptional programs broadly reflect aggressive disease behavior. **C.** Skin cutaneous melanoma (TCGA-SKCM) was the only cohort in which high cSCC-progression signature scores were associated with improved overall survival, highlighting divergence between melanoma and non-melanoma skin cancer progression programs. HNSC: head and neck squamous cell carcinoma, ACC: adrenocortical carcinoma, KIRC: kidney renal clear cell carcinoma, KIRP: kidney renal papillary cell carcinoma, LAML: acute myeloid leukemia, LGG: brain lower grade glioma, LIHC: liver hepatocellular carcinoma, LUAD: lung adenocarcinoma, MESO: mesothelioma, UVM: uveal melanoma, SKCM: skin cutaneous melanoma, OV: ovarian cancer.

Principal component analysis (PCA) analyses further demonstrated transcriptional separation between high- and low–score groups in these 12 TCGA cancer cohorts, in which the signature was correlated with overall survival (**Supplementary Figure S4**). These analyses support the notion that the cSCC-derived signature captures conserved tumor-progression programs across cancer types.

Collectively, these findings suggest that the molecular features underlying cSCC progression extend beyond cutaneous carcinoma and overlap with broader cancer-progression transcriptional programs.

### Identification of stage-associated candidate biomarkers for cSCC

We next sought to identify molecular biomarkers capable of complementing conventional clinical staging. To this end, each histological stage was systematically contrasted against all others in both HTG-Seq and GeoMx datasets. Genes demonstrating robust, consistent, and stage-discriminatory expression were retained (**Figure 5**). The corresponding log2 fold changes for the stage(s) in which they were differentially expressed are summarized in **Supplementary Table 3**.

**Figure 5.**
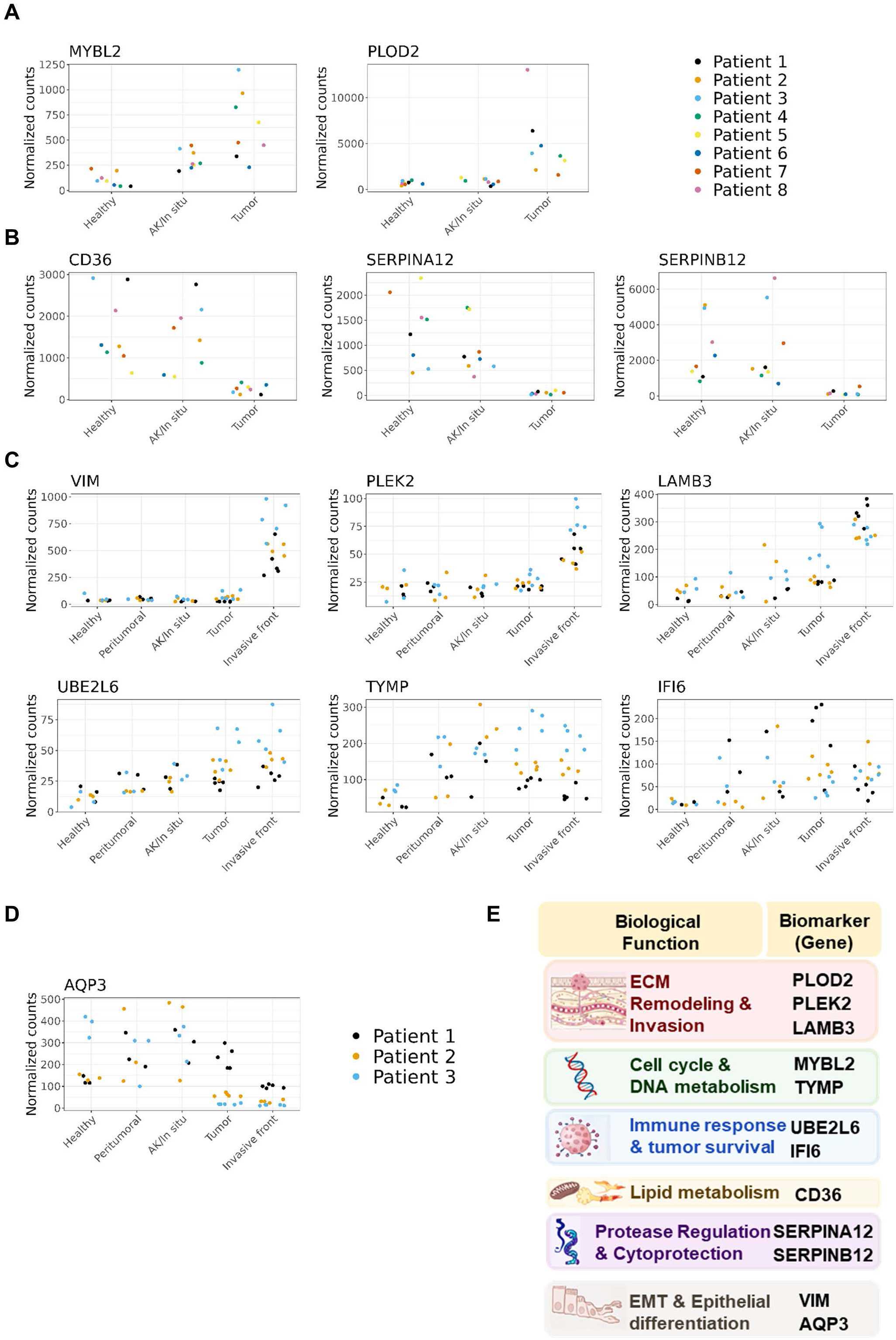
Identification of stage-associated candidate biomarkers for cSCC. **A-B.** Strip charts showing candidate biomarkers with progressive upregulation (**A**) or downregulation (**B**) across cSCC stages based on HTG-Seq bulk transcriptomic data. Upregulated candidates include *MYBL2* (MYB proto-oncogene like 2) and *PLOD2* (procollagen-lysine,2-oxoglutarate 5-dioxygenase 2), while downregulated candidates include *CD36* (fatty acid translocase), *SERPINA12* (serpin family A member 12), and *SERPINB12* (serpin family B member 12). **C-D.** Strip charts showing candidate biomarkers with progressive upregulation (**C**) or downregulation (**D**) across cSCC stages based on GeoMx spatial transcriptomic data. Upregulated candidates include *VIM* (vimentin), *PLEK2* (pleckstrin 2), *LAMB3* (laminin subunit beta 3), *UBE2L6* (ubiquitin-conjugating enzyme E2 L6), *TYMP* (thymidine phosphorylase), and *IFI6* (interferon alpha-inducible protein 6), while *AQP3* (aquaporin 3) is progressively downregulated. Biomarkers were selected based on robust stage vs rest expression patterns, and prioritized given the information of LRT analyses. **E.** Based on their biological functions, the selected biomarkers were organized into six categories: i) extracellular matrix remodeling and invasion (*PLOD2, PLEK2, LAMB3*), ii) cell cycle regulation and DNA metabolism (*MYBL2, TYMP,*), iii) immune activation and tumor cell survival (*UBE2L6, IFI6*), iv) lipid metabolism *(CD36)*, v) epithelial protease regulation (*SERPINA12* and *SERPINB12),* and vi) markers associated with invasive transition states (*VIM* and *AQP3)*.

Analysis of the HTG-Seq dataset identified *MYBL2* (MYB Proto-Oncogene Like 2) and *PLOD2* (Procollagen-Lysine,2-Oxoglutarate 5-Dioxygenase 2) as progressively upregulated during cSCC progression (**Figure 5A**). In contrast, *CD36* (Fatty Acid Translocase), *SERPINA12* (Serpin Family A Member 12), and *SERPINB12* (Serpin Family B Member 12) were consistently downregulated (**Figure 5B**). Parallel analysis of the GeoMx dataset revealed stage-dependent upregulation of *VIM* (Vimentin), *PLEK2* (Pleckstrin 2), *LAMB3* (Laminin Subunit Beta 3), *UBE2L6* (ubiquitin-Conjugating Enzyme E2 L6), *TYMP* (Thymidine Phosphorylase), and *IFI6* (Interferon Alpha-Inducible Protein 6) (**Figure 5C**), alongside progressive downregulation of *AQP3* (Aquaporin 3) (**Figure 5D**). Based on their biological function, the selected biomarkers were organized into six categories (**Figure 5E**).

To validate the spatial relevance of these candidates, we performed high-resolution spatial transcriptomic profiling using Visium HD, enabling direct visualization of gene expression within the tissue microenvironment and confirming stage-associated spatial patterns of selected biomarkers.

Consistent with HTG-Seq, *MYBL2* and *PLOD2* were selectively elevated in tumor regions and further increased at invasive stages (**Supplementary Figure S5A-B**), whereas *CD36*, *SERPINA12* and *SERPINB12* distinguished early (healthy/AK) from late (tumor) stages (**Supplementary Figure S5C-D**).

In agreement with GeoMx results, *VIM*, *PLEK2*, *LAMB3*, *UBE2L6*, *IFI6*, and *TYMP* exhibited stage-dependent upregulation, whereas AQP3 expression progressively declined (**Figure 6A-C, Supplementary Figure S6A-D**). Notably, Visium HD analyses revealed stage-restricted upregulation of *VIM* at the invasive front, accompanied by *AQP3* loss, defining a spatial invasiveness signature: AQP3⁺/VIM⁻ regions localized to tumor cores, whereas AQP3⁻/VIM⁺ regions marked invasive fronts (**Figure 6A, B**).

**Figure 6.**
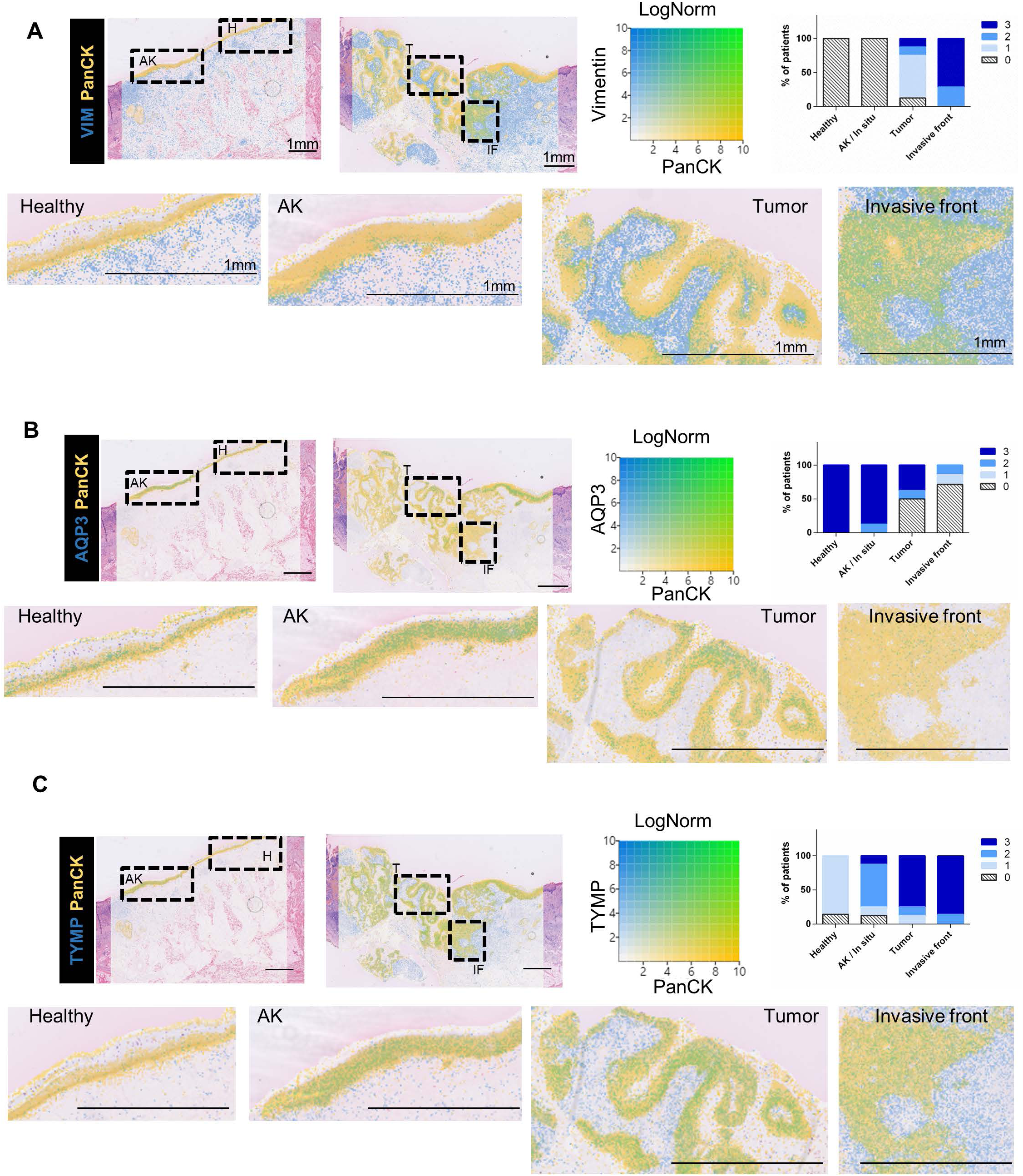
Dual AQP3 vs VIM spatial expression discriminates tumor core vs invasive front regions, while TYMP and IFI6 mark premalignant lesions. Cell-resolution spatial transcriptomic profiling was performed using 10x Genomics Visium HD on 8 skin samples spanning the full continuum of cSCC stages, from healthy tissue to invasive carcinoma. Representative Visium HD expression maps from a single patient are shown for *VIM*, *AQP3* and *TYMP*. **A.** Spatial visualization of *VIM* (blue) and *PanCK* (yellow) log-normalized expression generated using Loupe Browser, regions of co-expression are shown in green. VIM expression within PanCK⁺ epithelial regions is spatially restricted to the invasive front, consistent with localized activation of epithelial–mesenchymal transition–associated programs at sites of invasion. Scale bar, 1 mm. The right panel shows quantification of VIM expression intensity in PanCK⁺ regions across eight patients, scored from 0 (<5% positive area) to 3 (>60% positive area), and summarized as the percentage of patients per lesion type. **B.** Spatial visualization of *AQP3* (blue) and *PanCK* (yellow) expression demonstrating progressive loss of *AQP3* within PanCK⁺ epithelial regions across cSCC progression. The combined loss of *AQP3* and gain of *VIM* defines a dual spatial signature that robustly marks invasive tumor regions and enables early detection of invasion-associated states. Scale bar, 1 mm. Right panel shows AQP3 expression intensity scores (0 -3) in PanCK^+^ regions across eight patients. **C.** Spatial visualization of *TYMP* (blue) and PanCK (yellow) expression showing early upregulation of *TYMP* within PanCK⁺ epithelial regions in premalignant lesions, supporting potential utility of TYMP expression as biomarkers of high-risk premalignant and early disease states. Scale bar, 1 mm. Right panel shows *TYMP* expression intensity scores (0-3) in PanCK^+^ regions across eight patients.

Among early progression–associated biomarkers, TYMP was elevated from the AK/*in situ* stage onward in GeoMx analyses (see **Figure 5C**). Visium HD further confirmed low expression in healthy skin but marked upregulated in keratinocytes and tumor microenvironment components in diseased states (**Figure 6C**), supporting its potential prognostic relevance. Similarly, focal IFI6 “hot spots” were detected within selected AK lesions (**Supplementary Figure S6D**), suggesting a possible association with increased risk of progression, pending protein-level validation and functional studies.

Finally, investigation across 32 TCGA cohorts revealed that expression levels of *LAMB3*, *PLOD2*, and *SERPINB12* were significantly associated with overall survival in head and neck squamous cell carcinoma and multiple other cancer types (log-rank *P* < 0.05; **Supplementary Figure S7**), with directionality varying according to tumor context. These findings suggest that components of the cSCC progression program overlap with broader, yet context-dependent, tumor progression mechanisms operative across diverse malignancies.

## Discussion

We characterized the molecular evolution across the full continuum of cSCC progression by combining bulk transcriptomics (HTG-Seq) and spatial transcriptomics (GeoMx and Visium HD), using anatomically adjacent healthy skin, AK/cSCC *in situ,* invasive tumor core, and the invasive front. This within-field, multi-stage design addresses a key limitation of many prior cSCC transcriptomic studies, in which disease progression is inferred from cross-sectional comparisons across different patients ^16,17,19,20^. By anchoring multiple histological stages within a shared tissue context, our approach enables more accurate identification of progression-associated programs and facilitates nomination of candidate biomarkers that complement conventional clinical staging.

A central analytical strength of this study is the use of LRT to identify genes with statistically significant, stage-dependent expression trajectories. Unlike classical pairwise differential expression approaches, which test isolated contrasts between two stages, LRT modeling evaluates whether incorporating stage information improves model fit, enabling detection of monotonic as well as non-linear patterns across progression ^21,22^. This strategy is well suited for multistep processes such as cSCC evolution, where clinically relevant signals may emerge gradually or transiently rather than as binary on/off differences. Compared with stage-restricted or pairwise contrast designs commonly used in AK versus cSCC profiling ^23^, applying LRT also reduces the dependence on arbitrary stage-to-stage thresholds.

Consistent with a continuum model of cSCC, our data support a progressive shift in keratinocyte programs from differentiation-associated epidermal states toward proliferative, immune-interacting, and ECM-remodeling phenotypes, in agreement with prior bulk and single-cell/spatial studies ^16,17,19,24,25^. Within PanCK⁺ tumor-enriched regions, we identified distinct stage-linked programs, including interferon-stimulated and innate immune–associated signatures, alongside a separate ECM remodeling program that became more increasingly prominent in invasive tumor core and at the invasive front. These spatially patterned programs are consistent with prior transcriptomic studies of cSCC ^172627^ and with spatial atlas data showing tumor subpopulations and leading-edge niches ^19^. Together, these findings reinforce the concept that progression-associated biology is not uniformly distributed across lesions but is instead spatially organized into functionally specialized compartments.

Leveraging multi-stage pathway analyses, we nominated stage- and region-informative candidate biomarkers. Visium HD profiling revealed distinct localization patterns across the cSCC progression continuum, supporting their potential to delineate transitional or high-risk lesion states. Based on expression dynamics and biological function, biomarkers were grouped into three categories: genes upregulated in invasive cSCC, progressively downregulated genes, and biomarkers with reciprocal dynamics during invasive transitions (**Figure 7**). Invasion- and matrix remodeling-associated candidates were enriched in tumor and invasive regions, aligning with evidence linking these molecules to migration, invasion, EMT-associated programs, and laminin-332–linked aggressiveness in cSCC ^28–31^. Proliferation-associated biomarkers such as *MYBL2* increased with stage, in agreement with prior reports implicating *MYBL2* in cSCC progression ^32^. *TYMP* exhibited early and sustained upregulation, consistent with its documented expression in cutaneous tumors and its role in microenvironmental regulation ^33^. Interferon-related genes (*UBE2L6* and *IFI6*) increased from AK/*in situ* to invasive stages, aligning with studies linking interferon-inducible programs to keratinocytic lesion progression ^27,34^. Reciprocal *AQP3* and *VIM* expression distinguished tumor core from invasion-associated states, consistent with reports of VIM enrichment in metastatic primary cSCC ^35^ and lesion-specific, heterogeneous *AQP3* expression across keratinocyte neoplasia ^36,37^.

**Figure 7.**
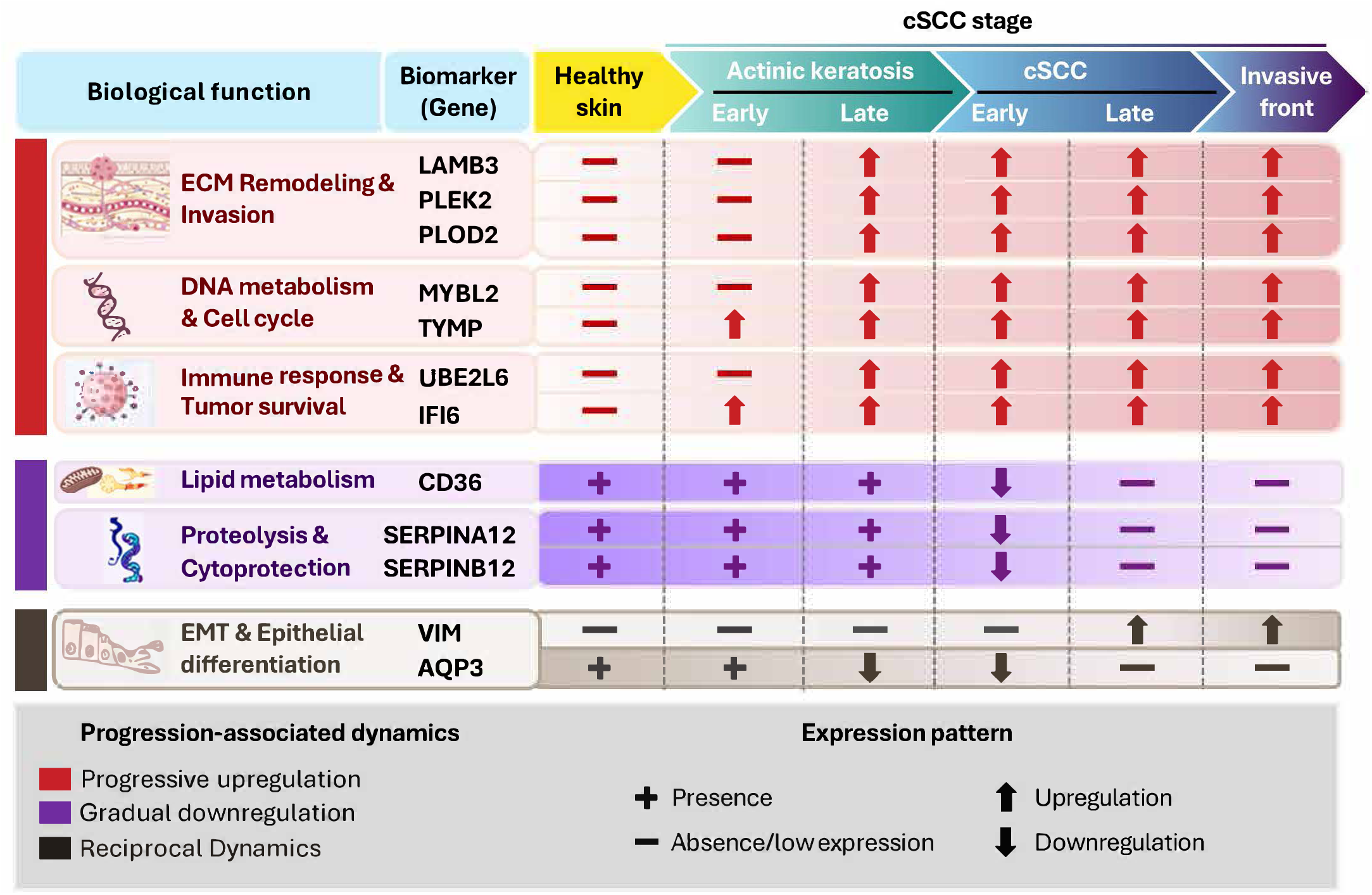
Stage-associated expression landscape of candidate biomarkers across cSCC progression. Schematic overview of biomarker expression patterns across six defined stages of cSCC, from healthy skin to invasive carcinoma. Biomarkers are organized into three predominant expression-pattern categories: (i) genes preferentially upregulated in invasive cSCC, including mediators of extracellular matrix remodeling and invasion (LAMB3, PLEK2, PLOD2), cell cycle regulation and DNA metabolism (TYMP, MYBL2), and immune activation and tumor pro-survival signaling (IFI6, UBE2L6); (ii) genes progressively downregulated during disease progression, including protease regulators and cytoprotective factors (SERPINA12, SERPINB12) and the lipid metabolism–associated receptor CD36; and (iii) markers associated with invasive transition states, including VIM and AQP3. Symbols (↑, ↓, +, −) denote relative expression trends or presence/absence across disease stages, providing an integrated framework for molecular staging and progression-associated phenotypes.

Regarding metabolism, our data indicate that metabolic reprogramming is compartment- and region-specific rather than uniform across lesions. While we previously reported substantial interpatient metabolic heterogeneity at each stage of cSCC progression ^38^, spatial profiling here demonstrates stage- and region-specific specific regulation within individual patients. Glycolysis and oxidative phosphorylation genes progressively declined in PanCK⁺ tumor cells but increased in CD68⁺ macrophages, consistent with prior descriptions of metabolic reprogramming in cSCC ^39–42^. Lipid metabolism pathways were among the most consistently downregulated programs, in agreement with bulk transcriptomic studies reporting reduced sphingolipid, triglyceride, and fatty acid–associated pathways in cSCC ^43^. Together, these data highlight compartment-specific metabolic adaptations and underscore the importance of spatial resolution for interpreting tumor metabolism.

A limitation of the present study is the relatively small sample size, reflecting the scarcity of specimens spanning the full continuum of cSCC progression. Notably, a previous meta-analysis required integration of 11 independent datasets and nonetheless lacked representation of invasive cSCC ^44^. To mitigate this limitation and evaluate the broader relevance of our findings, we extended our analysis to 32 TCGA cohorts comprising over 20,000 tumors, as previously described^45–48^. The results of our screening suggest that the identified cSCC-progression signature broadly reflects cancer progression at the RNA level across other 10 different cancer types.

Collectively, these results establish spatial transcriptomics as a powerful framework for modeling multistep carcinogenesis and provide a foundation for improving risk stratification, early detection, and precision management of patients with cSCC. Future studies integrating protein-level validation, longitudinal sampling, and functional analyses will be essential to translate these insights into clinically actionable strategies.

## Data availability

The authors declare that the data supporting the findings of this study are available within the paper and its supplementary information files. The datasets generated during the current study are accessible through the GEO accession number GSE319968 (GeoMx DSP) and GSE319969 (HTG-seq). The lists of genes found to be differentially expressed and the gene lists used for the heatmaps can be found in https://services.cbib.u-bordeaux.fr/cSCC_gene_tables/.

Additional data are available from the corresponding author upon reasonable request.

The R code used to reproduce the analyses will be made available upon publication at https://github.com/cbib/Reproduce_cSCC_GeoMx_HTGseq, and the TCGA Snakemake pipeline is available at https://github.com/cbib/snakemake_tcga_survival.

## Acknowledgments

The authors wish to thank Christine Alfaro (CHU Bordeaux, human sample collection), Dr. Olivier Cogrel, Dr. Jean-Michel Amici, Dr. Paul Cirotteau (Dermatopathology, CHU Bordeaux), Dr. Domitille Chalopin-Fillot (IBGC, Bordeaux University), Pr. Pierre Dubus, and Pr. Jean-Philippe Merlio (Tumor Bank and Tumor Biology Laboratory, Bordeaux University Hospital, Bordeaux, France).

High-throughput sequencing was performed by the ICGex NGS platform of the Institut Curie supported by the grants ANR-10-EQPX-03 (Equipex) and ANR-10-INBS-09-08 (France Génomique Consortium) from the Agence Nationale de la Recherche (“Investissements d’Avenir” program), by the ITMO-Cancer Aviesan (Plan Cancer III) and by the SiRIC-Curie program (SiRIC Grant INCa-DGOS-465 and INCa-DGOS-Inserm_12554). Data management, quality control and primary analysis were performed by the Bioinformatics platform of the Institut Curie.

