## Supplementary for "Spatial and Bulk Transcriptomic Profiling Defines the Molecular Evolution of Cutaneous Squamous Cell Carcinoma and Reveals Stage-Specific Biomarkers of Clinical Relevance": Naji et al_ supplementary data.docx

^1^ Univ. Bordeaux, Inserm, BRIC, UMR 1312, F-33076 Bordeaux, France.

^2^ Univ. Bordeaux, Centre de Bioinformatique de Bordeaux CBiB - US Ubx 006, F-33076 Bordeaux, France

^3^ Pathology Department, Bordeaux University Hospital, Bordeaux, France.

^4^ Univ. Bordeaux, CNRS, IBGC, UMR 5095, F-33000 Bordeaux, France

^5^ Explicyte Immuno-Oncology, Bordeaux, France

^6^ Institut Curie, PSL University, ICGex Next-Generation Sequencing Platform, 75005 Paris, France

^7^ Institut Curie, PSL University, Single Cell Initiative, 75005 Paris, France

^8^Dermatology Department, Bordeaux University Hospital, Saint-André Hospital, Bordeaux, France.

^#^ Authors contributed equally

^‡^ Joint supervision

**Running title:** Stage-resolved molecular programs in cSCC progression

^*^ **Corresponding Author**

Postal address: INSERM U1312, Bordeaux, F-33000 France

**Supplementary Data**

1- **Appendix S1-additional supplementary material file**
2- **Supplemental Tables (S1 to S4)**
3- **Supplemental Figures S1 to S7**

**Appendix S1-additional supplementary material file**

**Bulk RNA sequencing using HTG EdgeSeq RNA**

A total of 24 regions from eight Formalin-fixed, paraffin-embedded (FFPE) specimens were analyzed using the HTG EdgeSeq Transcriptome Panel (HTP; HTG Molecular Diagnostics, Tucson, AZ, USA) which employs a quantitative nuclease protection assay. The HTP includes 19,616 NanoString Probe Pairs (NPPs), comprising 19,398 target probes, 100 negative control probes, 92 external RNA control consortium (ERCC) RNA control probes, 22 genomic DNA probes, and 4 positive control probes (POS). The experimental workflow is illustrated in **Supplementary** **Figure S1A**.

**HTG EdgeSeq gene expression quantification**: Briefly, FFPE skin specimens were obtained from eight patients with cSCC. For each patient, tissue blocks contained histologically distinct regions corresponding to (i) healthy skin, (ii) actinic keratosis (AK) and/or *in situ* cSCC, and (iii) advanced invasive cSCC. Hematoxylin and eosin–stained sections were reviewed by a board-certified dermatopathologist (F.B.) to confirm diagnoses and guide region selection. Three regions of interest (healthy, AK/*in situ* cSCC, and advanced invasive cSCC) were macro-dissected separately from 5-µm unstained FFPE sections using annotated reference slides. Care was taken to minimize cross-contamination between regions. Sections underwent lysis and proteinase K digestion for 20 minutes at 56°C. Following protein inactivation, DNase treatment was performed at 37°C for 30 minutes. The quantitative nuclease protection assay was then carried out using the HTG EdgeSeq Processor according to the manufacturer’s instructions. Adapters and sample-specific barcodes were added during PCR amplification to generate sequencing-ready libraries. The resulting libraries were sequenced on an Illumina NextSeq 2000 platform (P2 Flow cell, 100 cycles). The resulting FASTQ files were processed towards a gene expression count matrix using the HTG EdgeSeq Reveal Software. The quality control (QC) of NGS data was assessed prior analysis to ensure the data robustness.

**GeoMx Digital Spatial Profiling**

Spatial transcriptomics was performed using the Nanostring GeoMx Digital Spatial Profiler (GeoMx) ^1^. The experimental workflow is illustrated in Supplementary **Figure S1B**. The protocol consists of five main steps: (i) standard FFPE tissue preparation; (ii) incubation with a mixture of visualization markers (VMs) and DSP probes; (iii) imaging and region of interest (ROI) selection; (iv) ultraviolet (UV) photocleavage; and (v) oligonucleotide collection and quantification using the Nanostring nCounter® system.

**Tissue Preparation and FFPE processing:** Formalin-fixed, paraffin-embedded (FFPE) tissue sections (eight unstained, 5 µm thick) were mounted on charged slides, and one adjacent section was stained with hematoxylin and eosin (H&E). The H&E slide was scanned at 20x and 40x magnification using the 3DHISTECH slide scanner and reviewed by a board-certified pathologist (F.B.) to delineate tumor and adjacent normal regions. Four unstained slides were processed for Opal Multiplex immunofluorescence (IF) staining (PerkinElmer), and two unstained sections were used for Digital Spatial Profiling (DSP) at Explicyte Immuno-Oncology (Bordeaux, France). Multispectral IF staining was performed using the Opal four-color IHC Kit (NEL794001KT; Perkin Elmer, Waltham, MA, USA) according to the manufacturer’s instructions. This system employs tyramide signal amplification (TSA)-conjugated fluorophores to enhance target detection. Briefly, the slides were deparaffinized in xylene, rehydrated through graded ethanol solutions (100%, 95%, 70% for 2 min) and washed in Tris-buffered saline containing 0.1% Tween-20 (TBST, Merck). Heat-induced epitope retrieval was perfomed by microwave treatment (45 seconds at full power followed by 15 minutes at 20% power) in antigen retrieval Buffer. Sections were blocked with blocking diluent (Perkin Elmer) for 10 minutes at room temperature. Primary antibodies were incubated for 37°C for 32 minutes using a Dako Slide Hybridizer (Agilent technologies, CA, USA). After washing in TBST, slides underwent additional microwave treatment in antigen retrieval buffer prior to sequential incubation with subsequent antibodies. Slides were then counterstained with DAPI for 5 minutes, mounted with ProLong Gold antifade reagent (Invitrogen) and coverslipped. Imaging was performed using the Vectra 3.0 spectral imaging system (PerkinElmer), and image analysis was conducted with inForm image software (PerkinElmer). Whole-slide low-magnification scans (10x) were first acquired to ensure alignment with pathologist-annotated H&E sections.

In total, 43 to 46 areas of illumination (AOIs) were then selected across tumor cores, immune-infiltrated areas, invasive fronts, and stromal compartments (**supplementary Figure S2-S3, Supplementary Table S2**) to capture spatial heterogeneity within each tissue section.

**Selection of region of interests and gene expression profiling:** Briefly, tissue sections were stained with a CD68/PanCK/Vimentin/Sytox panel to visualize tissue architecture and ROIs selection. For each slide, ROIs were drawn in order to analyze the expression of the WTA gene panel in the healthy regions, AK/cSCC in situ, center of cSCC, and invasive front. Each ROI was further segmented into three to five area of illumination (AOIs) based on marker expression. In total, 43 to 46 AOIs were selected across tumor cores, immune-infiltrated areas, invasive fronts, and stromal compartments (**supplementary Figure S2-S3, Supplementary Table S2**) to capture spatial heterogeneity within each tissue section. PanCK-enriched AOIs represented epithelial/tumor regions; CD68-enriched AOIs captured macrophage-rich compartments; and PanCK⁺/Vimentin⁺ AOIs were used as an operational proxy for the invasive front, consistent with histopathologic evalutation.

Gene expression of each AOI was profiled using the GeoMx Whole Transcriptome Atlas (WTA) panel, which interrogates 18,695 protein-coding genes. Slides were incubated overnight with *in situ* hybridization (ISH) probes conjugated to unique DNA indexing oligonucleotides via a UV-photocleavable linker. Each AOI was then sequentially illuminated with a focused UV laser to release the corresponding DNA barcodes, which were collected into a 96-well plate. GeoMx aspirates collected in 96-well plates contained photocleaved DNA oligonucleotide comprising an analyte identifier, a unique molecular identifier (UMI) barcode, and primer binding sites. PCR amplification was performed to incorporate Illumina adapter sequences and unique dual sample indices. Following a pooling purification, the final library was sequenced on an Illumina Novaseq6000 platform using a dual-index workflow.

Quality control (QC) of NGS data was assessed prior to downstream analysis to ensure data robustness and to exclude low-quality AOIs. Different parameters were analyzed and notably the raw read threshold, alignment, and sequencing saturation. AOIs with fewer than 1,000 raw reads, less than 80% aligned reads, or less than 50% sequencing saturation were excluded from further analysis. Moreover, segments with exceptionally low signal (< 1% of panel genes detected above the limit of quantification, based on negative control probes) were removed.

**GeoMx and HTG EdgeSeq analyses**

**Quality control:** An unsupervised clustering of the normalized expression data was performed to identify potential outliers which could affect the data analysis. Principal component analysis (PCA) and hierarchical clustering (heatmap visualization) were conducted prior to differential expression analysis. One patient profiled with GeoMx was excluded from the analyses due to a pronounced batch effect. This patient was found out to be different from the rest of the samples, likely because it was processed in a different batch and AOIs were larger than the rest of patients. No additional outliers were detected.

**Differential expression analysis:** For each cellular compartment, count matrices were analyzed independently using DESeq2. For each celluar compartment (PanCK+ and CD68+), gene-level differential expression across disease stages was assessed using DESeq2 likelihood ratio tests (LRT), comparing a full model including stage (~ tissue) to a reduced intercept-only model (~1). This approach identifies genes whose expression varies significantly across multiple disease stages without assuming a specific pairwise contrast.

**Gene expression pattern clustering:** Genes with a Benjamini–Hochberg adjusted *P* < 0.01 were considered significantly differentially expressed and retained for downstream analyses. Significantly differentially expressed genes were subjected to rlog transformation and clustered using DEGreport to identify dominant expression patterns across tissue compartments. Cluster ordering was adjusted for visual clarity, and smoothed expression trajectories were plotted using locally estimated scatterplot smoothing (LOESS) regression.

**Gene set export and pathway enrichment analyses:** For each expression cluster, gene lists were extracted and exported as structured tables for pathway analysis. Functional enrichment analyses were performed using g:Profiler (via gprofiler2 R package), followed by pathway redundancy reduction and summarization using the ORSUM Python package, as detailed in the Supplementary Pathway Analyses folder on GitHub (see Code availability).

**Pathway selection for heatmap visualization:** To facilitate interpretation and visualization of enrichment results, ORSUM-filtered Gene Ontology (GO: Biological Process) outputs were manually summarized for heatmap display. For each gene-expression trend, top-ranked GO terms were reviewed and grouped when representing closely related biological processes. Genes associated with these terms were retrieved from the enrichment results and merged into a single, non-redundant gene set for each pathway category. This procedure was applied exclusively for visualization purposes to ensure comprehensive representation of dominant biological signals while minimizing redundancy, without modifying the underlying enrichment analyses or statistical results.

**Stage-specific biomarker identification:** Keratinocyte-enriched (PanCK⁺) count matrices were analyzed to identify candidate biomarkers capable of discriminating stages of cSCC progression. Differential expression analyses were performed using DESeq2 Wald tests, in which each histological compartment was systematically compared against all other compartments. Genes showing statistically significant differential expression after multiple-testing correction (adjusted *P* < 0.05, Benjamini Hochberg correction), sufficient expression differences (log2FC > 1.5 or log2FC < -1.5), and only found in a specific stage were retained for further evaluation.

Candidates genes were then prioritized based on (i) effect size, (ii) cross-patient reproducibility (consistent direction of change across patients), and (iii) spatial concordance between GeoMx and Visium HD datasets. Finally, biological plausibility was assessed by evaluating consistency with prior literature on keratinocyte differentiation, EMT/invasion, immune interaction, and cSCC progression. Literature support was used as contextual evidence rather than as a statistical selection criterion.

**Visium HD Spatial transcriptomics**

**Spatial transcriptomic sample preparation:** The spatial gene expression process including probe hybridization, probe Ligation, enabled probe release with CytAssist and extension was performed using Visium CytAssist Spatial Gene Expression for FFPE, Human Transcriptome, 6.5mm, 4 rxns (PN-1000675, 10x Genomics) according to the manufacturer’s instructions. Briefly, tissue sections were decrosslinked at 80°C for 30 minutes. Human whole transcriptome pairs of probes targeting 18000 genes were hybridised overnight. Paired probes were ligated and transferred onto the Visium slide using the CytAssist system.

The spatial gene libraries were genertated using probe-based library construction (10x Genomics). Library quantification and quality assessment were performed using Qubit fluorometric assay (Invitrogen; dsDNA HS Assay Kit), Agilent 2100 Bioanalyzer (High Sensitivity DNA chip) and KAPA Library Quantification Kit for Illumina platforms (Roche). Libraries were sequenced on an NovaSeq 6000 (Illumina) using paired-end 28 x 50 bp reads at a sequencing depth of 25,000 reads per spot.

**Spatial transcriptomic data processing**: SpaceRanger software v2.0.0 (10x Genomics) was used for processing spatial transcriptomic data. The raw base call (BCL) files were demultiplexed and mapped to the reference genome. Loupe Browser (10x Genomics) was used to align the barcoded spot patterns to histological images, perform tissue selection and conduct the pathological annotation.

**Evaluation of stage-specific biomarkers using Visium HD:** Spatial transcriptomic Visium HD (8 µm resolution) datasets were analyzed using Loupe Browser v9 (10x Genomics) to evaluate spatial expression of selected candidate biomarkers (see GeoMx section). For each gene, spatial expression was visualized in the “Features” panel and displayed as a co-expression map with an epithelial reference signature (“PanCK”), defined by KRT5, KRT6A, KRT8, KRT17, and KRT19, to localize expression within epithelial and tumor compartments and ensure alignment with GeoMx PanCK⁺ regions. Expression values were displayed using Loupe’s log-normalized UMI scale, normalized to total UMI counts per spot. For visualization, PanCK expression is shown in yellow, biomarker expression in blue, and co-expression in green. To facilitate semi-quantitative comparison across histological regions and patients, biomarker signal intensity was scored per region as follows: 0 (<5% positive area), 1 (5–10%), 2 (10–20%), and 3 (>20% positive area). Semi-quantitative assessment was performed using ImageJ (NIH) following color-channel separation, threshold adjustment, area measurement, and logical co-localization analysis using the Image Calculator function.

**Snakemake - The Cancer Genome Atlas survival analysis pipeline**

To enable systematic and reproducible survival analyses across multiple The Cancer Genome Atlas (TCGA) cohorts, we developed a modular computational pipeline implemented in Snakemake. The workflow was designed to integrate transcriptomic and clinical data, performs gene- and signature-level expression stratification, and evaluates associations with overall survival in a standardized manner across cancer types. Briefly, the pipeline performs the following steps: (1) automatic retrieval of TCGA RNA-seq cohorts from the Genomic Data Commons (GDC) data portal; (2) mapping of Ensembl identifiers to gene symbols using BioMart; (3) cohort-specific normalization of gene expression using DESeq2’s median-of-ratios method; (4) stratification of patients into extreme expression percentiles based on individual gene expression (for single-gene analyses) or single-sample gene set enrichment analysis (ssGSEA) scores (for multi-gene analyses); (5) generation of Kaplan–Meier survival curves and PCA visualizations; and (6) production of corresponding statistical summary tables for downstream interpretation. Notably, our Snakemake pipeline also enables integration of gene sets annotated as “upregulated” and “downregulated” by computing a combined score in which downregulated ssGSEA scores are subtracted from upregulated scores, as previously described ^2^, providing a robust and flexible framework for leveraging differentially expressed gene signatures derived from in-house or published datasets.

***Data sources and preprocessing:*** Bulk RNA sequencing data and associated clinical annotations were obtained from TCGA using the TCGAbiolinks R package, which downloads the updated datasets via the Genomics Data Portal (GDC). Ensembl gene identifiers were converted to HGNC gene symbols via biomaRt R package. Gene-level raw counts generated from STAR alignments were imported into DESeq2 and normalized using the median-of-ratios method to account for library size and compositional bias. Normalized expression values were used for downstream analyses.

Clinical data were extracted from TCGA harmonized metadata and included vital status, days to death, and days to last follow-up. For censored cases (patients alive at last follow-up), missing values for days to death were replaced by days to last follow-up, as in the TCGAbiolinks R package. Overall survival time was expressed in years for visualization purposes.

***Gene signature definition:*** Gene signatures were provided as a tab-delimited input file, with each column representing a distinct signature and containing comma-separated HGNC gene symbols. Signatures were classified into two categories:

1. **Single-gene signatures**, containing one gene only.
2. **Multi-gene signatures**, consisting of two or more genes.

For multi-gene signatures, enrichment scores were computed using single-sample gene set enrichment analysis (ssGSEA). When signatures were provided as matched *_UP* and *_DOWN* pairs sharing a common base name, a combined score was derived by subtracting the *_DOWN* score from the *_UP* score, thereby generating a single composite signature.

***Single-sample gene set enrichment analysis:*** ssGSEA was performed via the GSVA R package on normalized gene expression values. Enrichment scores were calculated independently for each patient and each multi-gene signature, using normalized enrichment and default kernel density estimation parameters. The resulting score matrix was transposed to represent patients as rows and signatures as columns.

For single-gene signatures, normalized expression values of the corresponding gene were directly used as patient-level scores.

***Patient stratification:*** To assess robustness to stratification choice, we evaluated multiple percentile cut-offs (25%, 33%, 50%). For non-median cutoffs, intermediate samples were excluded to focus on extreme expression groups; median split retained all patients. For each threshold, patients with scores below the lower percentile were classified as Low, those above the corresponding upper percentile as High, and those in between as Intermediate.

Only patients classified as Low or High were retained for survival analyses, while intermediate cases were excluded to maximize contrast between groups. Categorical stratification variables were stored alongside clinical metadata for reproducibility.

***Survival analysis:*** Overall survival was assessed using Kaplan–Meier estimators implemented in the survival and survminer R packages. Survival objects were constructed using time-to-event data (in years) and vital status, with death treated as the event of interest.

For each gene signature and percentile cut-off, survival curves were compared between High and Low expression groups using the log-rank test. Hazard ratios (HRs) and corresponding 95% confidence intervals (CIs) were calculated using a Cox proportional hazards regression model with group (High vs. Low) as the predictor variable. The Low group was used as the reference category. Confidence intervals were derived from the Wald statistic of the Cox model.

Kaplan–Meier plots were generated only for comparisons meeting the significance threshold (*P* < 0.05). Plots included group-specific sample sizes in the legend and were exported at publication-quality resolution.

***Principal component analysis:*** For signature–percentile combinations reaching survival significance threshold, principal component analysis (PCA) was performed as an exploratory visualization. Variance-stabilized expression values were computed using the DESeq2 variance-stabilizing transformation. PCA was conducted on the subset of patients included in the survival analysis (Low and High groups only), and samples were coloured according to expression group.

The first two principal components were plotted, with the percentage of explained variance indicated on each axis. PCA figures were exported at high resolution.

***Output generation and reproducibility:*** For each TCGA project, the pipeline generated the following outputs:

1. A matrix of patient-level signature scores.
2. A table of categorical patient stratifications across all signatures and percentile cutoffs.
3. A complete matrix of log-rank *P* values.
4. A filtered *P* value table retaining only statistically significant associations.

All analyses were executed in R, within a Snakemake-controlled environment to ensure reproducibility, traceability of parameters, and consistency across TCGA projects.

**Supplemental Tables**

**Table S1. Clinical characteristics of the patients included in HTG-Seq and GeoMx.**

| **Patient Number** | **HTG-Seq** | **GEOMX** | **Sex** | **Age** | **Location** | **Immune status** | **Histopathological subtypes** |
| --- | --- | --- | --- | --- | --- | --- | --- |
| **1** | **+** | **+** | **M** | **77** | **Scalp** | **KTR** | **Invasive, moderately differentiated cSCC** |
| **2** | **+** | **+** | **M** | **88** | **Scalp** | **IC** | **Invasive, moderately to poorly differentiated cSCC with neurotropism** |
| **3** | **+** | **+** | **M** | **84** | **Upper limb** | **IC** | **Invasive, moderately differentiated cSCC** |
| **4** | **+** |  | **M** | **68** | **Scalp** | **HTR** | **Invasive, poorly differentiated cSCC** |
| **5** | **+** | **-** | **M** | **83** | **Left arm** | **IC** | **Invasive, moderately differentiated cSCC** |
| **6** | **+** |  | **M** | **92** | **Scalp** | **IC** | **Invasive, poorly differentiated cSCC with neurotropism** |
| **7** | **+** |  | **M** | **94** | **Vertex / Scalp** | **IC** | **Invasive, moderately differentiated, keratinizing cSCC** |
| **8** | **+** |  | **M** | **95** | **Scalp** | **KTR** | **Invasive, poorly differentiated cSCC with neurotropism** |

IC, Immunocompetent; KTR, Kidney transplant recipient; HTR, Heart Transplant Recipient.

### **Table S2. Distribution of areas of illumination (AOIs) across regions of interest (ROI) and patients in the GeoMx analysis.** Summary of the number of AOIs generated per region of interest (ROI) and per patient in the GeoMx Digital Spatial Profiler experiment. Tissue sections were stained with a CD68/PanCK/Vimentin/Sytox multiplex panel to visualize tissue architecture and guide ROI selection. For each slide, ROIs corresponding to healthy skin, peritumoral regions, AK/ in situ CSCC, invasive cSCC core, and invasive front were defined. Each ROI was subsequently segmented into three or five AOIs based on PanCK, Vimentin, and CD68 expression to enable cell-type–resolved profiling using the Whole Transcriptome Atlas (WTA) panel.

| Patient Number | Tissue | Segment | Number of AOIs |
| --- | --- | --- | --- |
| 1 | Healthy | Macrophages | 3 |
|  |  | PanCK | 3 |
|  | AK/*In situ* | Macrophages | 3 |
|  |  | PanCK | 3 |
|  | Invasive front | Macrophages | 5 |
|  |  | PanCK | 5 |
|  |  | PanCK.Vimentin | 5 |
|  | Peritumoral | Macrophages | 3 |
|  |  | PanCK | 3 |
|  | Tumor | Macrophages | 5 |
|  |  | PanCK | 5 |
| 2 | Healthy | Macrophages | 3 |
|  |  | PanCK | 3 |
|  | AK/*In situ* | Macrophages | 3 |
|  |  | PanCK | 3 |
|  | Invasive front | Macrophages | 5 |
|  |  | PanCK | 5 |
|  |  | PanCK.Vimentin | 5 |
|  | Peritumoral | Macrophages | 3 |
|  |  | PanCK | 3 |
|  | Tumor | Macrophages | 5 |
|  |  | PanCK | 5 |
| 3 | Healthy | Macrophages | 3 |
|  |  | PanCK | 3 |
|  | AK/*In situ* | Macrophages | 3 |
|  |  | PanCK | 3 |
|  | Invasive front | Macrophages | 5 |
|  |  | PanCK | 5 |
|  |  | PanCK.Vimentin | 5 |
|  | Peritumoral | Macrophages | 3 |
|  |  | PanCK | 3 |
|  | Tumor | Macrophages | 5 |
|  |  | PanCK | 5 |
| 4 (Excluded) | Healthy | Macrophages | 5 |
|  |  | PanCK | 4 |
|  | AK/*In situ* | Macrophages | 3 |
|  |  | PanCK | 3 |
|  | Invasive front | Macrophages | 5 |
|  |  | PanCK | 5 |
|  |  | PanCK.Vimentin | 5 |
|  | Peritumoral | Macrophages | 3 |
|  |  | PanCK | 3 |
|  | Tumor | Macrophages | 5 |
|  |  | PanCK | 5 |

**Table 3. Log2 fold changes of candidate biomarkers identified by stage-wise differential expression analysis in HTG-Seq and GeoMx datasets**

| **HTG-Seq markers** | | | | | | | | |  |  |
| --- | --- | --- | --- | --- | --- | --- | --- | --- | --- | --- |
|  | Healthy vs others | | Healthy vs AK/*in situ* | | Healthy vs Tumor | | Ak/*in situ* vs Tumor | |  |  |
| **Gene symbol** | log2FC | padj | log2FC | padj | log2FC | padj | log2FC | padj |  |  |
| **MYBL2** | -2.15 | 1.79E-05 | 1.52 | 7.43E-04 | 2.62 | 1.00E-200 | 1.11 | 7.85E-03 |  |  |
|  | Tumor vs others | | Healthy vs AK/*in situ* | | Healthy vs Tumor | | Ak/*in situ* vs Tumor | |  |  |
| **Gene symbol** | log2FC | padj | log2FC | padj | log2FC | padj | log2FC | padj |  |  |
| **PLOD2** | 2.59 | 5.44E-17 | 0.43 | 5.62E-01 | 2.59 | 1.00E-200 | 2.16 | 1.00E-200 |  |  |
| **CD36** | -2.68 | 9.63E-16 | 2.44E-03 | 9.98E-01 | -2.57 | 1.00E-200 | -2.57 | 1.00E-200 |  |  |
| **SERPINA12** | -4.58 | 1.37E-34 | -0.50 | 4.92E-01 | -4.81 | 1.00E-200 | -4.30 | 1.00E-200 |  |  |
| **SERPINB12** | -3.92 | 3.12E-20 | 0.08 | 9.51E-01 | -3.93 | 1.00E-200 | -4.01 | 1.00E-200 |  |  |
| **GeoMx DSP markers** | | | | | | | | | | |
|  | Invasive Front vs others | | Healthy vs peritumor | | Healthy vs AK/*in situ* | | Healthy vs Tumor | | Healthy vs  Invasive front | |
| **Gene symbol** | log2FC | padj | log2FC | padj | log2FC | padj | log2FC | padj | log2FC | padj |
| **VIM** | 3.56 | 2.00E-78 | 0.23 | 8.23E-01 | -0.06 | 9.50E-01 | 0.36 | 2.78E-01 | 3.76 | 1.00E-200 |
| **PLEK2** | 1.57 | 1.32E-33 | 0.19 | 8.94E-01 | 0.22 | 8.00E-01 | 0.60 | 3.68E-02 | 1.91 | 2.12E-09 |
| **LAMB3** | 1.76 | 2.86E-10 | 0.20 | 8.98E-01 | 0.91 | 3.76E-01 | 1.70 | 4.58E-06 | 3.01 | 1.00E-200 |
| **AQP3** | -2.08 | 3.60E-09 | 0.35 | 7.90E-01 | 0.70 | 2.96E-01 | -1.47 | 1.90E-02 | -2.42 | 5.34E-07 |
|  | Healthy vs others | | Healthy vs  Invasive front IF | | Peritumor vs Invasive front | | AK/*in situ* vs Invasive front | | Tumor vs  Invasive front | |
| **Gene symbol** | log2FC | padj | log2FC | padj | log2FC | padj | log2FC | padj | log2FC | padj |
| **UBE2L6** | -1.52 | 1.29E-06 | 2.00 | 9.30E-07 | 0.99 | 7.22E-04 | 0.64 | 1.90E-02 | 0.19 | 3.30E-01 |
| **TYMP** | -1.62 | 4.69E-09 | 1.40 | 1.08E-04 | -0.13 | 7.95E-01 | -0.58 | 1.27E-01 | -0.38 | 5.29E-02 |
| **IFI6** | -2.56 | 4.89E-10 | 2.34 | 1.88E-08 | 1.03 | 1.01E-01 | 0.09 | 8.95E-01 | -0.30 | 5.16E-01 |

**Supplementary Figures and figure legends**

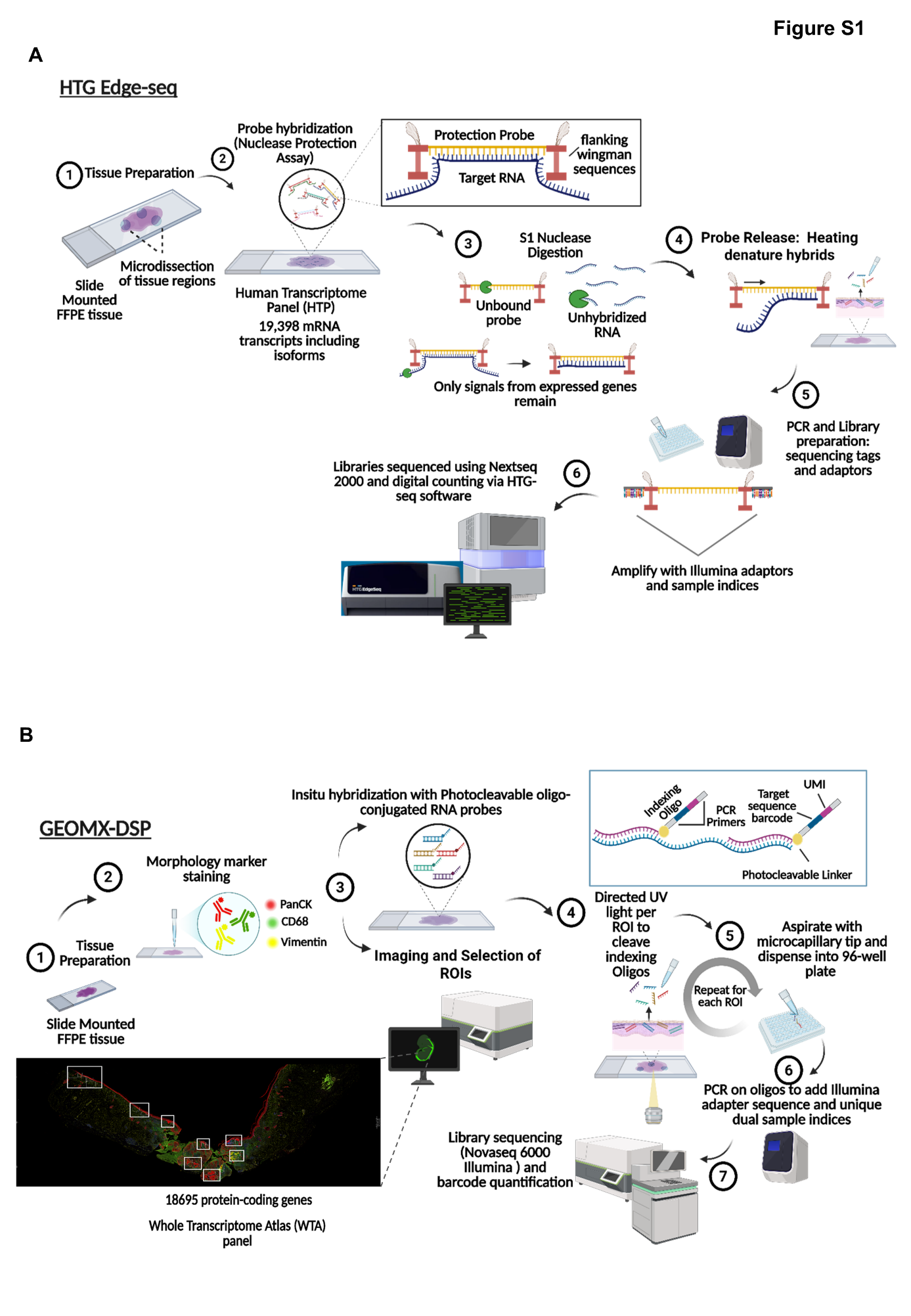

**Figure S1. Experimental workflow of HTG EdgeSeq and GeoMx.**

**A.** Technical overview of the workflow for HTG EdgeSeq. FFPE tissue sections were prepared using standard protocols, followed by microdissection of regions representing distinct lesion stages. Gene-specific probe hybridization was performed, followed by tissue lysis and proteinase K treatment, DNase digestion, and release of protected probes. PCR was used to add sequencing adapters and sample indices, and libraries were sequenced on an Illumina NextSeq 2000 (P2 flow cell, 100 cycles). Resulting FASTQ files were processed using HTG EdgeSeq Reveal Software to generate gene expression count matrices. This platform uses a targeted nuclease protection assay in which gene-specific probes hybridize to RNA transcripts, protecting them from nuclease degradation. Sequencing of the protected probe fragments after heat denaturation enables quantitative bulk gene expression profiling without spatial resolution.

**B.** Technical workflow for GeoMx Digital Spatial Profiler (DSP). FFPE tissue sections were incubated with a multiplexed visualization marker panel (CD68, PanCK, Vimentin, and Sytox) to define tissue architecture and guide region of interest (ROI) selection. Following incubation with DSP probes, ROIs were exposed to ultraviolet (UV) light to release indexing oligonucleotides, which were collected for each area of illumination (AOI). Oligos were PCR-amplified to add Illumina adapters and unique dual indices, and libraries were sequenced on an Illumina NovaSeq 6000. Gene expression for each AOI was quantified using the GeoMx Digital Spatial Profiler (NanoString®) Whole Transcriptome Atlas (WTA) panel. Created with Biorender

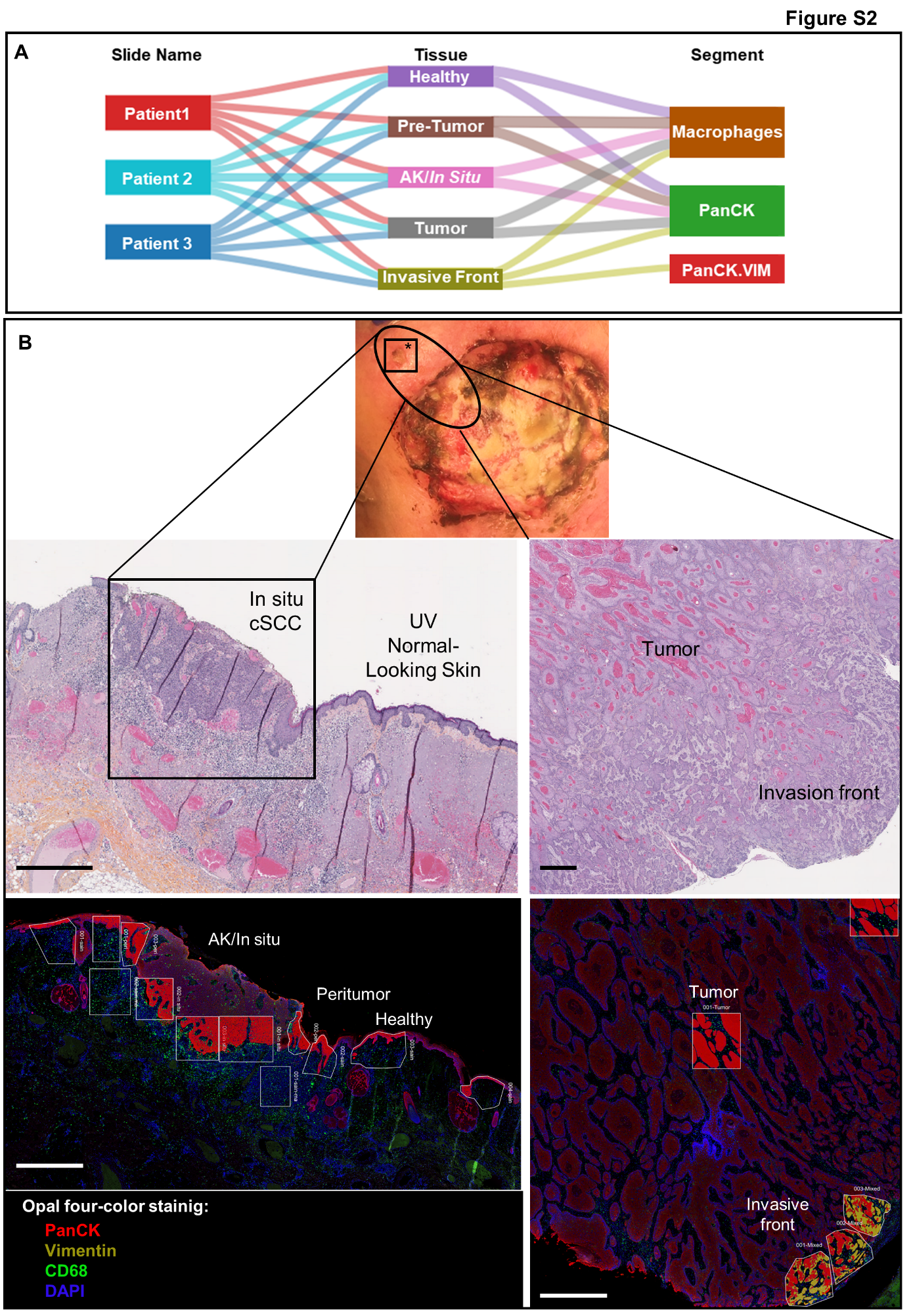

**Figure S2. GeoMx study design and multi-stage sampling strategy**

**A**. Panel A summarizes the multi-stage ROI selection strategy applied across three independent patients.

**B.** Representative overview of the GeoMx workflow. A patient presenting with *in situ* cSCC and invasive cSCC within the same lesion field is shown as an example. Hematoxylin and eosin (H&E) staining illustrates the identification of multiple histological stages within a single biopsy, including adjacent healthy skin, peritumoral region, cSCC *in situ*, invasive tumor core, and invasive front.

Tissue sections were subsequently stained using a multiplex immunofluorescence panel (CD68, PanCK, Vimentin, Sytox) to visualize tissue architecture and guide region-of-interest (ROI) selection for GeoMx profiling. CD68⁺ cells correspond to macrophages, PanCK⁺ regions to epithelial/tumor compartments, and PanCK⁺/Vimentin⁺ areas to the invasive tumor front. Scale bar, 1 mm.

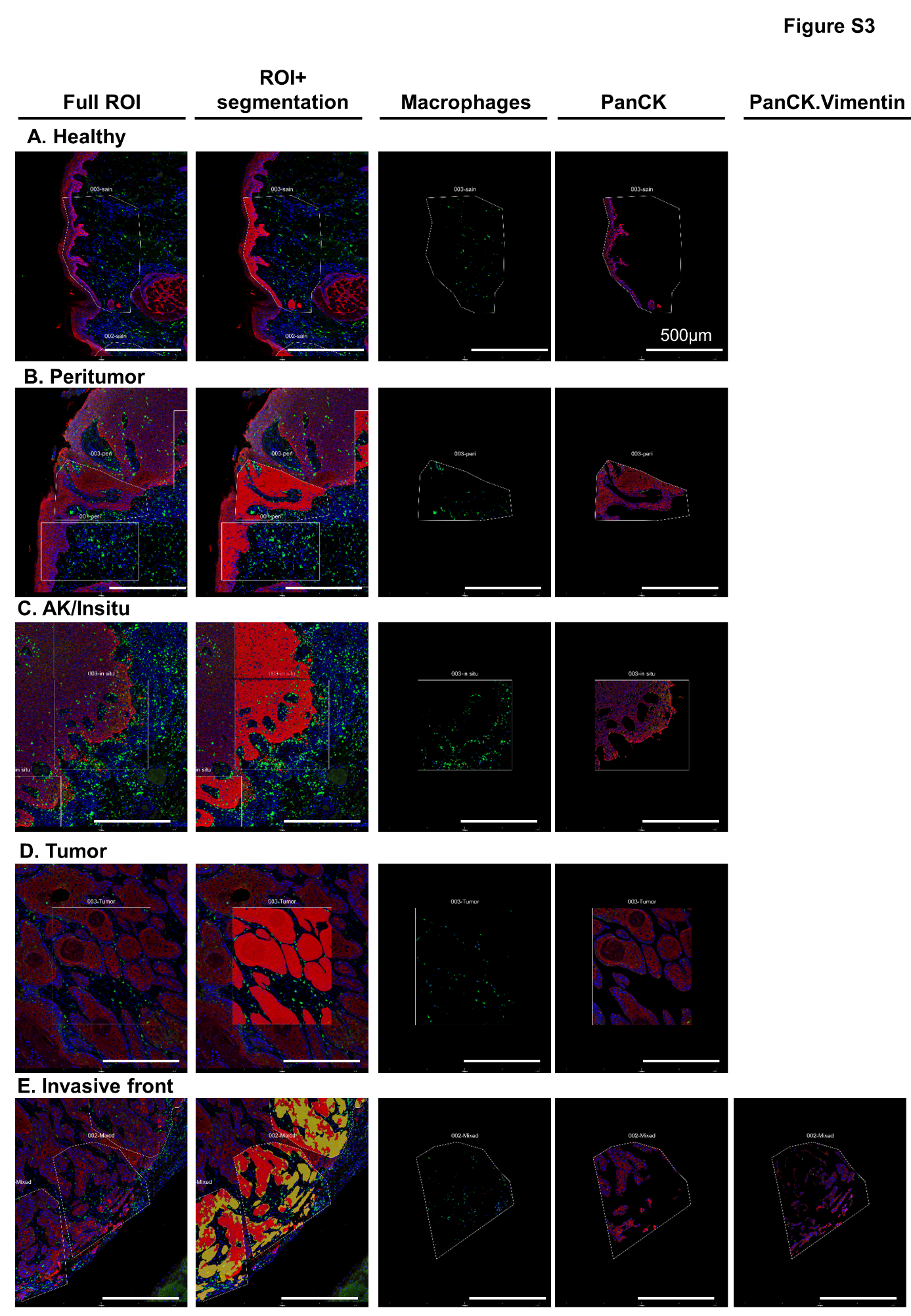

**Figure S3. ROI segmentation and AOI definition for GeoMx profiling**

Skin biopsies were obtained from four patients who exhibited a continuous histopathological spectrum within a single tissue section, including adjacent healthy skin, AK/ cSCC *in situ*, invasive cSCC core, and the invasive front. For each stage, representative examples are shown, including the full ROI, segmented compartments, and corresponding areas of illumination (AOIs). ROIs were segmented into AOIs based on multiplex staining for CD68, PanCK, and Vimentin, enabling compartment-specific transcriptomic profiling. AOIs included macrophage-enriched (CD68⁺), epithelial/tumor (PanCK⁺), and invasive front–associated (PanCK⁺/Vimentin⁺) regions. Scale bar, 500 µm.

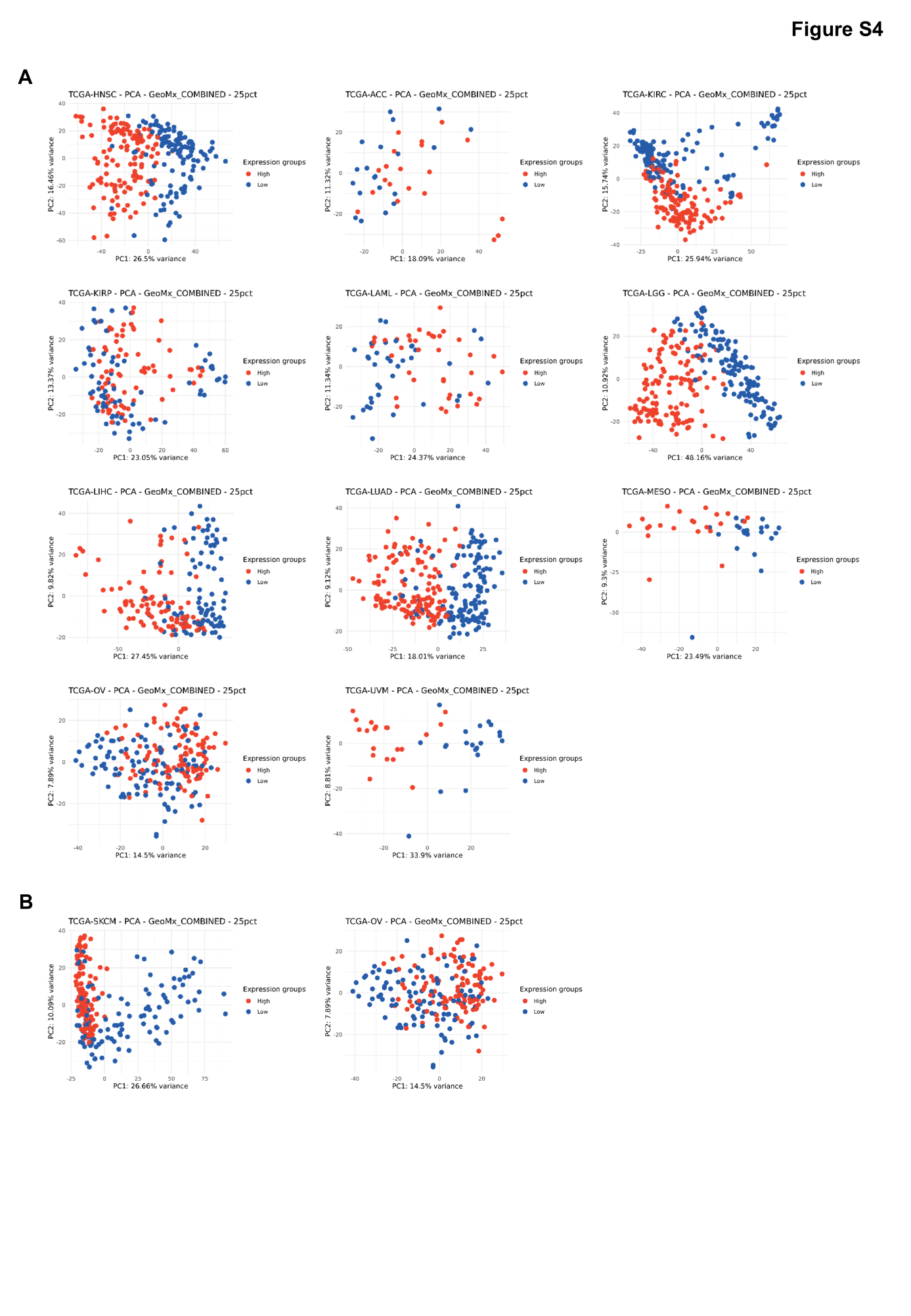

**Figure S4. Principal component analysis of TCGA cohorts stratified by the cSCC-progression signature.**

Principal component analysis (PCA) of TCGA cohorts for which the cSCC-progression signature showed a significant association with overall survival (log-rank *P* < 0.05). PCA was performed on variance-stabilized gene expression values, after DESeq2 normalization, from patients included in the survival analysis. Samples are colored according to progression-score group. Axes represent the first two principal components, with the percentage of variance explained indicated. These PCA plots provide an unsupervised visualization of phenotypic transcriptional separation associated with the cSCC-progression signature across cancer types.

**A.** PCA plots of the 10 cohorts in which the cSCC-progression signature correlated with poor overall survival. From left to right and from top to bottom: adrenocortical carcinoma (TCGA-ACC), cervical squamous cell carcinoma and endocervical adenocarcinoma (TCGA-CESC), head and neck squamous cell carcinoma (TCGA-HNSC), kidney renal clear cell carcinoma (TCGA-KIRC), kidney renal papillary cell carcinoma (TCGA-KIRP), acute myeloid leukemia (TCGA-LAML), brain lower grade glioma (TCGA-LGG), liver hepatocellular carcinoma (TCGA-LIHC), lung adenocarcinoma (TCGA-LUAD), mesothelioma (TCGA-MESO), and uveal melanoma (TCGA-UVM).

**B.** PCA plot of metastatic cutaneous melanoma (TCGA-SKCM), which was the only cohort in which cSCC-progression signature correlated with good overall survival.

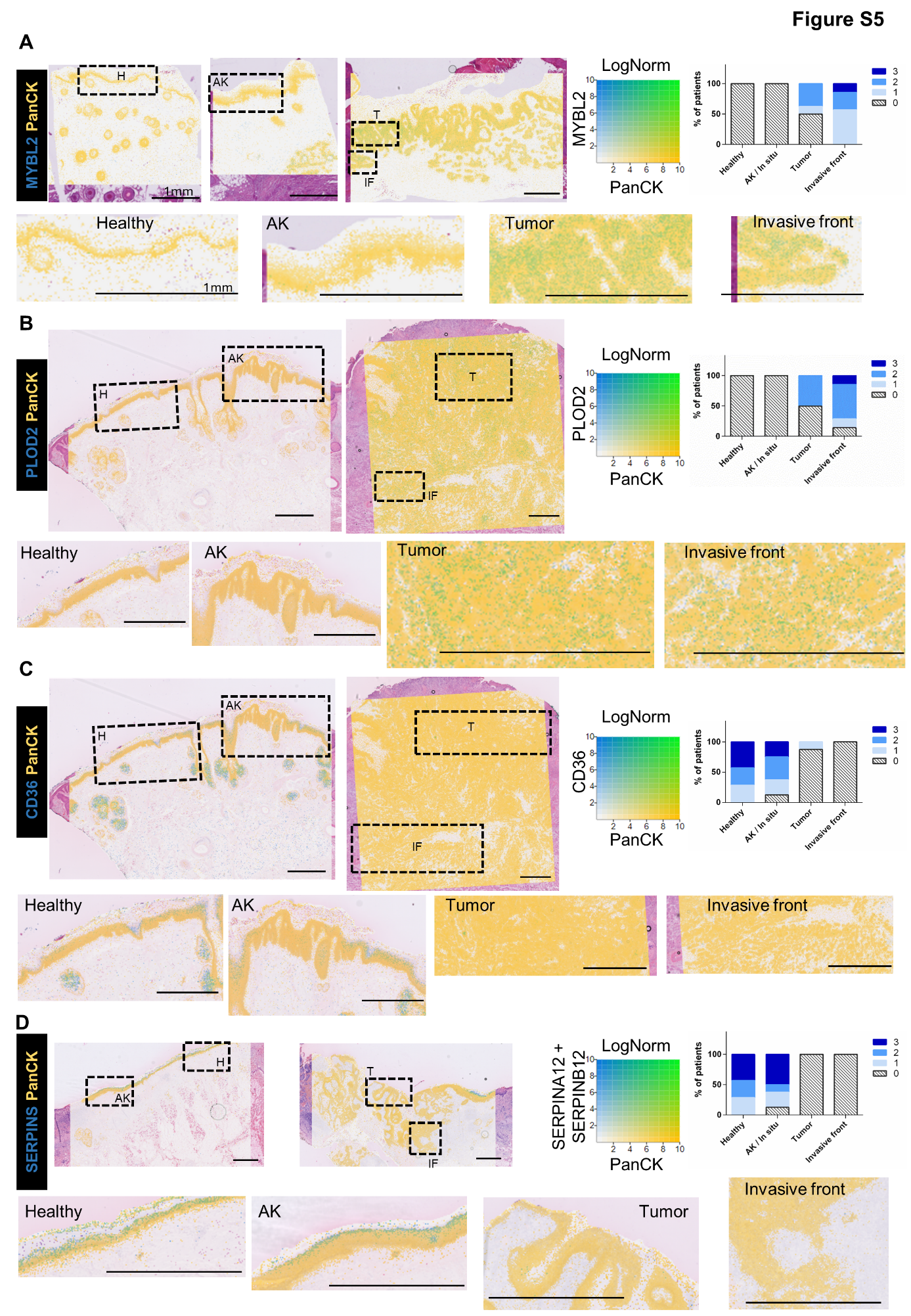

**Figure S5.** **Spatial visualization of stage-associated expression of *MYBL2*, *PLOD2*, *CD36*, *SERPINA12*, and *SERPINB12* across cSCC progression**

High-resolution spatial transcriptomic profiling was performed using 10x Genomics Visium HD on tissue sections containing healthy skin, carcinoma *in situ*, and invasive cSCC.

**A–D**. Loupe Browser visualization of *MYBL2*/*PanCK* (**A**), *PLOD2*/*PanCK* (**B**), *CD36*/*PanCK* (**C**), and *SERPINA12* & *SERPINB12*/*PanCK* (**D**) expression. Co-expression is shown in green. Scale bar, 1 mm. Right panel shows RNA expression intensity scores (0-3) in PanCK^+^ regions across eight patients.

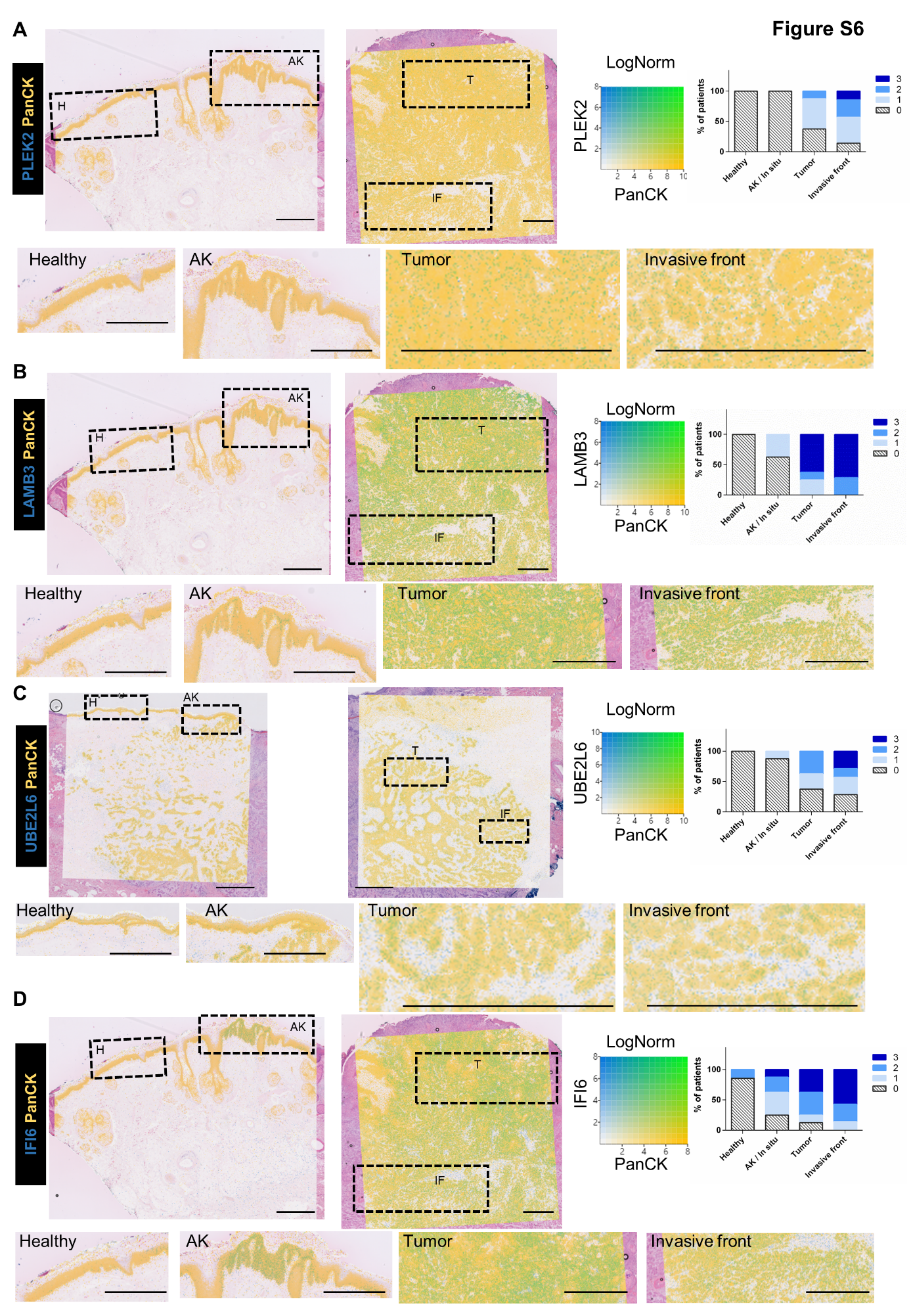

**Figure S6.** **Spatial visualization of stage-associated expression of *PLEK2*, *LAMB3*, *UBE2L6*, and *IFI6* across cSCC progression.**

High-resolution spatial transcriptomic profiling was performed using 10x Genomics Visium HD on skin samples spanning the full cSCC continuum, from healthy tissue to invasive carcinoma.

**A–D**. Loupe Browser visualization of *PLEK2*/*PanCK* (**A**), *LAMB3*/*PanCK* (**B**), *UBE2L6*/*PanCK* (**C**), and *IFI6*/*PanCK* (**D**) expression. Co-expression is shown in green. Scale bar, 1 mm. Right panel shows *RNA* expression intensity scores (0-3) in PanCK^+^ regions across eight patients.

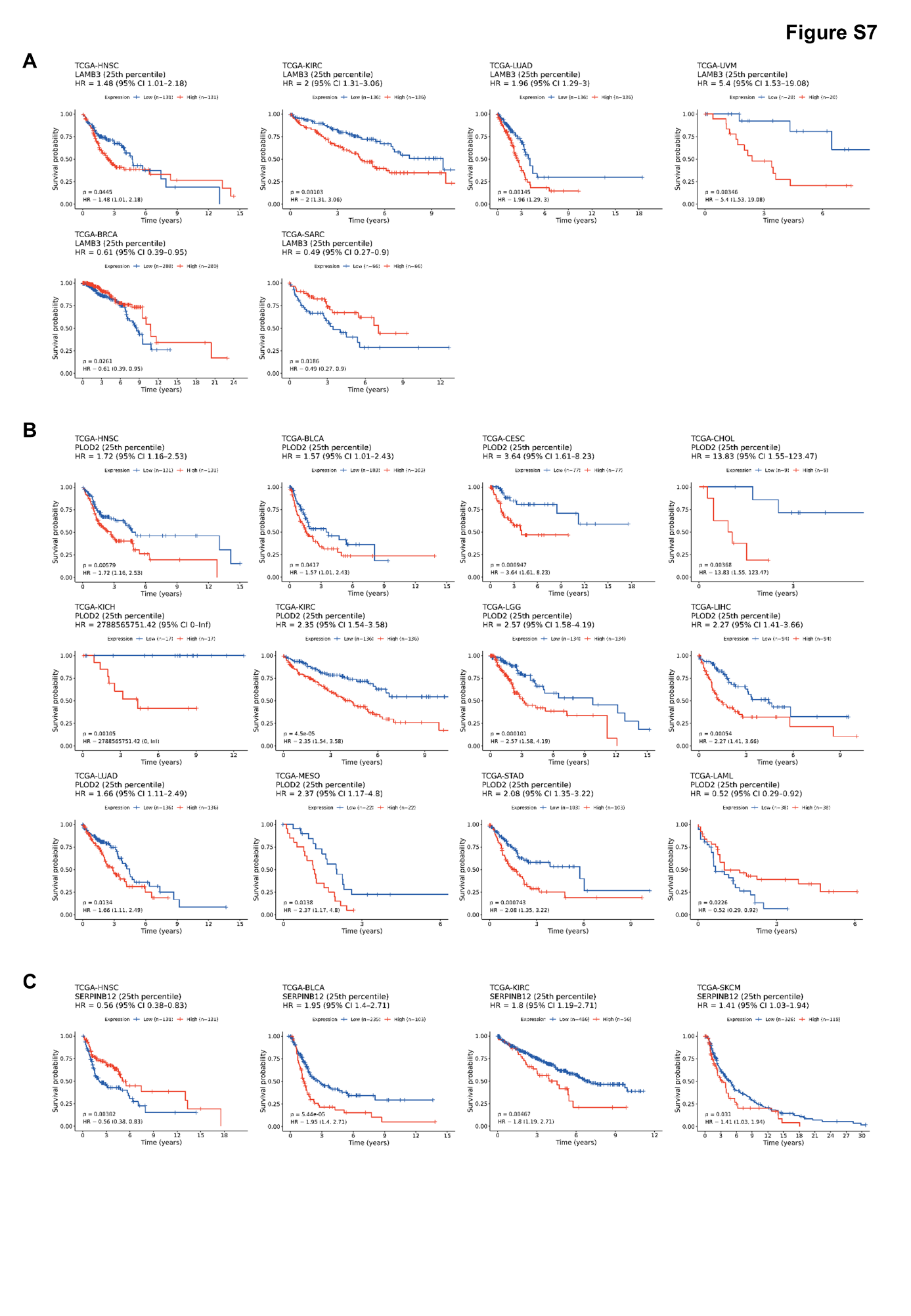

**Figure S7. Association of *LAMB3*, *PLOD2*, and *SERPINB12* expression with overall survival across TCGA pan-cancer cohorts.**

Kaplan–Meier survival analyses were performed across 32 TCGA cancer types to evaluate the association between gene expression and overall survival. Patients were stratified according to gene expression levels, and log-rank tests were used to assess statistical significance (P < 0.05).

**A–C**. Association between expression of *LAMB3* (**A**), **PLOD2** (*B*), and *SERPINB12* (**C**) and overall survival across TCGA cohorts. Expression of each gene was significantly correlated with overall survival in multiple cancer types.

Cancer type abbreviations: HNSC, head and neck squamous cell carcinoma; KIRC, kidney renal clear cell carcinoma; LUAD, lung adenocarcinoma; UVM, uveal melanoma; BRCA, breast invasive carcinoma; SARC, sarcoma; BLCA, urothelial bladder carcinoma; CESC, cervical and endocervical cancers; CHOL, cholangiocarcinoma; KICH, kidney chromophobe; KIRP, kidney renal papillary cell carcinoma; LGG, lower grade glioma; LIHC, liver hepatocellular carcinoma; MESO, mesothelioma; STAD, stomach adenocarcinoma; LAML; acute myeloid leukemia; SKCM, skin cutaneous melanoma.
